# Metabolic collaboration between cells in the tumor microenvironment has a negligible effect on tumor growth

**DOI:** 10.1101/2022.02.08.479584

**Authors:** Johan Gustafsson, Fariba Roshanzamir, Anders Hagnestål, Sagar M. Patel, Oseeyi I. Daudu, Donald F. Becker, Jonathan L. Robinson, Jens Nielsen

**Affiliations:** Department of Biology and Biological Engineering, Chalmers University of Technology, Gothenburg, Sweden; Wallenberg Center for Protein Research, Chalmers University of Technology, Gothenburg, Sweden; Hagnesia AB, Hindås, Sweden; Department of Biochemistry and Redox Biology Center, University of Nebraska-Lincoln, Lincoln, Nebraska 68588, United States; BioInnovation Institute, Copenhagen, Denmark

## Abstract

The tumor microenvironment is comprised of a complex mixture of different cell types interacting under conditions of nutrient deprivation, but the metabolism therein is not fully understood due to difficulties in measuring metabolic fluxes and exchange of metabolites between different cell types *in vivo*. Genome-scale metabolic modeling enables estimation of such exchange fluxes as well as an opportunity to gain insight into the metabolic behavior of individual cell types. Here, we estimated the availability of nutrients and oxygen within the tumor microenvironment using concentration measurements from blood together with a metabolite diffusion model. In addition, we developed an approach to efficiently apply enzyme usage constraints in a comprehensive metabolic model of human cells. The combined modeling reproduced severe hypoxic conditions and the Warburg effect, and we found that limitations in enzymatic capacity contribute to cancer cells’ preferential use of glutamine as a substrate to the citric acid cycle. Furthermore, we investigated the common hypothesis that some stromal cells are exploited by cancer cells to produce metabolites useful for the cancer cells. We identified over 200 potential metabolites that could support collaboration between cancer cells and cancer associated fibroblasts, but when limiting to metabolites previously identified to participate in such collaboration, no growth advantage was observed. Our work highlights the importance of enzymatic capacity limitations for cell behaviors and exemplifies the utility of enzyme constrained models for accurate prediction of metabolism in cells and tumor microenvironments.

## Introduction

The tumor microenvironment (TME) consists of many different cell types that share a common influx of metabolites from the blood, and its leaky blood vessels make diffusion the main mechanism for transport of metabolites and oxygen from the blood to the cells^1–4^. Furthermore, the blood vessel density is unevenly distributed, leading to a highly variable availability of nutrients and oxygen, where hypoxic and even necrotic regions tend to be more common in the center of the tumor^5^. Cancer cells residing in the TME are generally thought to strive for proliferation, and the metabolic behavior that is optimal for maximizing growth will vary with the availability of metabolites. Known such behaviors are lactate export (the Warburg effect^6^), which is shared with other highly proliferative cells such as expanding T cells^7^, and “glutamine addiction”^8^ where the tricarboxylic acid (TCA) cycle is preferentially fed with glutamine. Furthermore, bypass of mitochondrial complex I can be used to increase ATP production at a lower substrate efficiency in cells with high ATP demand^9^, although such a behavior has not explicitly been proven to exist in cancers.

Collaboration scenarios between different “healthy” cell types and tumor cells, where cells such as fibroblasts and macrophages are thought to be exploited by the cancer cells, has been a topic of great interest for the last decade^10–13^. Specifically, these cells are thought to regulate the tumor microenvironment through signaling as well as provide nutrients such as lactate and pyruvate to the tumor cells to enhance their growth. Partial evidence of the latter has been presented, where cancer-associated fibroblasts have been observed to secrete metabolites that were taken up by cancer cells^12^, but there is no solid evidence that such collaboration is directly beneficial for growth.

The study of collaborative metabolism in tumors is difficult for two reasons: 1) It is challenging to create a realistic *in vitro* representation of the tumor microenvironment using cell lines. For example, the influx of oxygen and all other metabolites from blood needs to be controlled and constant over time; and 2) It is difficult to measure metabolite uptake rates *in vivo* for all metabolites, although isotope labeling has successfully been employed for single metabolites^14^. An alternative approach is therefore needed.

Genome-scale modeling^15^ of human metabolism using flux balance analysis (FBA) involves performing in silico analyses of a reaction network under steady-state conditions and has been used to investigate metabolism in for example muscles^9^, tumors^16^, and Alzheimer’s disease^17^. A recent approach is the GECKO framework^18^, which enables the integration of enzyme kinetic data with genome-scale metabolic models to simulate more physiologically meaningful flux distributions even when metabolite exchange rate data is limited.

In this project, we developed a diffusion model for constraining metabolite uptake rates in tumors and applied enzyme usage constraints to a human genome-scale model to simulate tumor metabolism. Our metabolic models showed that glucose and oxygen are the most limiting substances for growth. Furthermore, our models predicted glutamine addiction, where fueling the TCA cycle with glutamine can yield more ATP since it requires less enzyme usage per ATP produced. In addition, we observed that metabolic collaboration between cancer-associated fibroblasts (CAFs) and cancer cells likely has a very small effect on tumor growth.

## Results

### Simulation of tumor cell growth

We used the genome-scale metabolic model Human1^19^ enhanced with enzyme constraints to model the metabolism of cancer cells in the tumor microenvironment. The model was manually curated (Methods) and a non-growth associated maintenance (NGAM) requirement was added as an ATP cost of 1.833 mmol gDW^−1^ h^−1^ derived from literature^20,21^. We developed a lightweight version of the GECKO toolbox^18,22^, called GECKO Light (Note S1), which similar to GECKO constrains the total metabolic enzyme usage based on k_cat_ values from the BRENDA database^23^. The main improvement in GECKO Light is a substantially decreased execution time (~2 minutes on a standard laptop computer) and a smaller generated model size.

We assumed that diffusion is the dominant mechanism for influx of metabolites into tumors^24^ and developed a diffusion model that uses the metabolite concentrations in blood and their diffusion coefficients to estimate the metabolite uptake constraints (Fig. 1A, Note S2). Instead of estimating absolute uptake constraints, which are expected to vary across cells depending on many factors such as distance to capillaries, we estimated relative uptake constraints according to

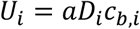

where *U*_*i*_ is the estimated upper bound for the uptake flux of metabolite *i, a* is a proportionality constant that is inversely related to the distance to the closest capillaries, *D*_*i*_ is the diffusion coefficient for metabolite *i*, and *c*_*b,i*_ is the concentration of metabolite *i* in the blood. While the value of *a* varies across cells and is difficult to estimate for a given cell, its value will be the same for all metabolites within the same cell, which makes it possible to investigate the metabolism at different pseudodistances from blood vessels. In addition, we developed a blood flow model, where the blood flow into the tumor rather than diffusion was assumed to be the limiting factor for delivery of metabolites to cells and the influx was modeled to be directly proportional to the contents of the blood (Note S3).

**Fig. 1:**
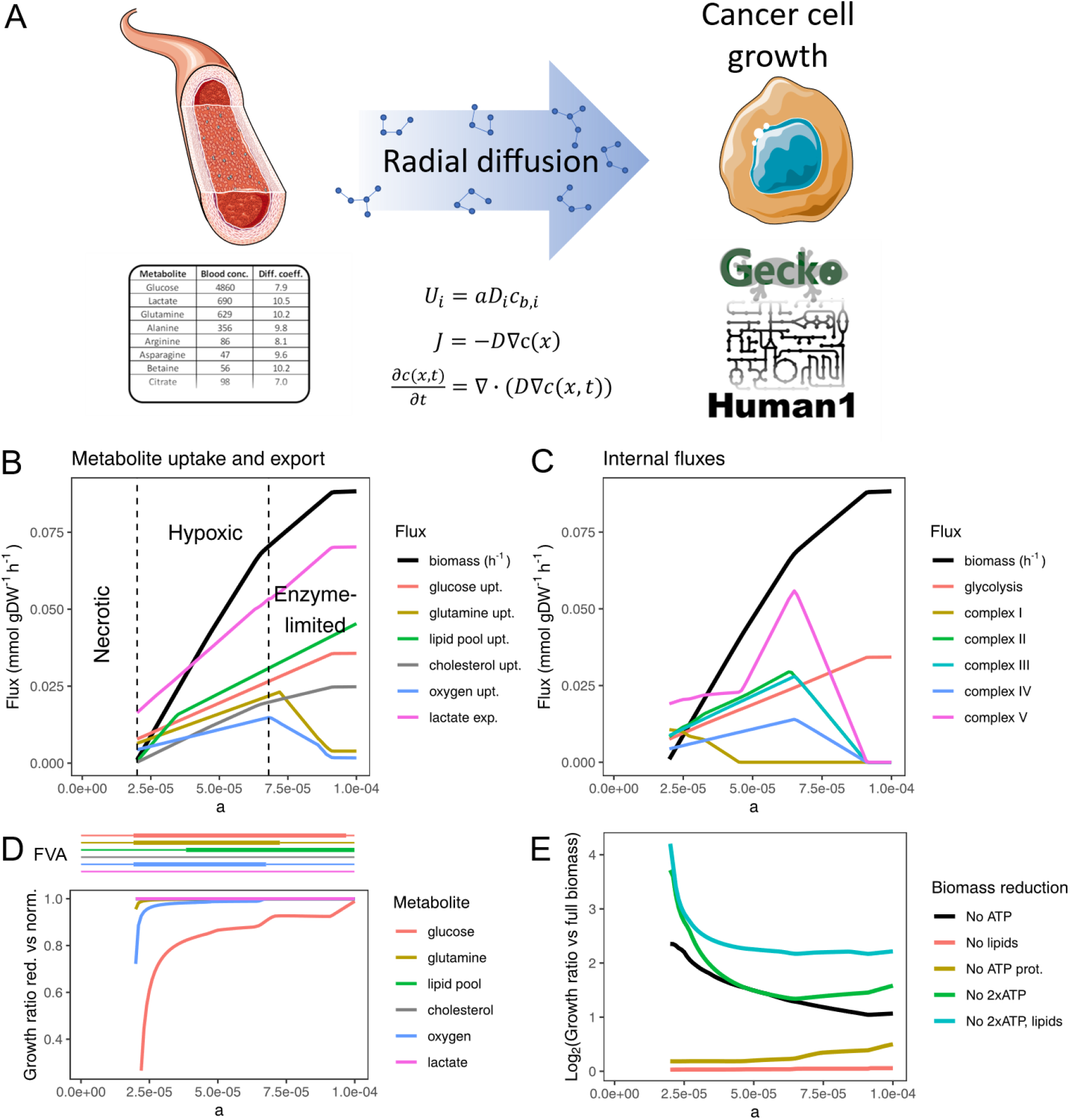
Modeling cancer cell growth in the tumor microenvironment. A. Modeling setup. The metabolite uptake bounds are estimated from a diffusion model based on metabolite concentrations in blood and their diffusion coefficients. In addition, enzyme usage constraints are added to the model using GECKO Light. B. Simulated specific growth rate (biomass production) and metabolite exchange rates for different values of a using the diffusion model. Some fluxes are scaled to enable visualization of all metabolites in the same figure: glucose upt.: 0.01, glutamine upt.: 0.05, Lipid pool upt.: 0.2, cholesterol upt.: 10, oxygen upt.: 0.1, and lactate exp.: 0.01. C. Internal fluxes in the same simulations as B. The glycolytic flux is multiplied by 0.01 and the fluxes through the complexes are multiplied by 0.1. D. Growth dependence of metabolites. (Top) A thick line indicates a required uptake rate of at least 95% of a metabolite for maintaining growth, as determined by flux variability analysis (FVA). (Bottom) The simulated specific growth rate of the model is compared to that when the maximum uptake rate of a single metabolite is reduced to 90%. The effect from lipid pool, cholesterol, and lactate is negligible or non-existent, although some of them are identified as limiting by the FVA. E. Change in specific growth rate when removing parts of the biomass reaction. The figure shows the specific growth rate ratio between models with reduced and original biomass reaction. In all cases, the model is optimized for growth. “No ATP prot.” refers to removal of the ATP cost from polymerizing amino acids into proteins, while “No 2xATP” refers to having both the protein generation ATP cost and the growth-associated maintenance (GAM) ATP cost removed from the biomass reaction. For “No 2xATP, lipids”, the consumption of lipids have also been removed in addition to the other two factors. ATP generation is the limiting factor for all values of a. With ATP costs removed, the availability of lipids became limiting. See Fig. S5 for details.

We retrieved blood plasma measurements from several sources to form a collection of 69 metabolites with associated concentration values^25–28^ (Note S4; Table S1, S2, and S3; Fig. S1). Lipids were grouped into sterols (represented by cholesterol) and other lipids (fatty acyls, glycerolipids, glycerophospholipids, and sphingolipids, represented as a mix of fatty acids). For oxygen we used the concentration of free oxygen, which can diffuse, excluding roughly 98% of the total concentration in blood that is bound to hemoglobin.

We collected 18 metabolite diffusion coefficients from literature^29–31^, and estimated the remaining coefficients using a linear model based on molecular mass^32^ (Fig. S2, Note S5, Table S3). Lipids are not soluble in water and are therefore transported together with albumin or a lipoprotein, or potentially as droplets. These are large particles that do not diffuse well, and we modeled the diffusion of these particles by using the diffusion coefficient of albumin for all lipids.

We first simulated the tumor cell growth for different values of the proportionality constant *a* using FBA (Fig. 1B). For small values of *a* (greater distances from capillaries), the model cannot produce enough ATP to uphold cell maintenance requirements, resulting in necrosis. For higher values of *a*, we see a hypoxic region where the cells can survive and grow but are limited in growth by availability of nutrients and oxygen, followed by a region where the enzyme constraints become the major limiting factor for growth. At small values of *a*, the limited oxygen supply was primarily used for oxidative phosphorylation, which provided much ATP per substrate. However, when more nutrients were available a larger specific growth rate could be reached by allocating more of the enzymatic capacity to glycolysis instead of to these large and slow enzyme complexes. The simulation predicts an early bypass of complex I, while the remaining electron transport chain (ETC) complexes remain active at a wider *a* range (Fig. 1C), which is consistent with a previous study where complex I bypass was described to increase ATP yield at the cost of a lower substrate efficiency^9^. The model exhibits an extreme behavior where oxidative phosphorylation is completely shut down for large values of *a*. In practice, such a behavior is generally not observed. 80% of the ATP has for example been reported to be generated by oxidative phosphorylation in some highly proliferative cells^33^, but regardless of this limitation the model can still serve as a means to understand observed metabolic behaviors.

Next, we evaluated which metabolites were limiting for growth (Fig. 1D, S3). Glucose and oxygen were by far the two most important metabolites for growth, followed by a smaller effect from glutamine, while the other limiting metabolites had a small effect (Fig. 1D). Neither lactate nor albumin was limiting for growth since neither of them can give a positive ATP contribution without using oxygen. For the hypoxic range, free access to oxygen yielded a clear increase in growth (Fig. S4). To investigate which cellular processes are limiting for growth we repeated our simulations with different parts of the biomass reaction removed, which showed that ATP production was the main limiting factor (Fig. 1E, S5). To assess the sensitivity of our results, we repeated our analyses with the growth-associated ATP cost and NGAM reduced to 50% and 25% of their original values, which confirmed that glucose and oxygen were still the most important contributors to growth (Fig. S6). The blood flow model produced similar results, although with a substantially smaller hypoxic region (Fig. S7). The major difference is a higher availability of lipids and oxygen, which increases oxygen usage at small *a* values.

### Amino acid metabolism

The diffusion model predicted large uptake fluxes of glutamine, glycine, serine, and threonine, and export of proline and aspartate, while the rest of the amino acids were consumed at lower fluxes, primarily for protein synthesis (Fig. 2A, Fig. S8). The model did not predict glutamate secretion, which is observed in some cancer cell lines^34,35^. While glutamate export has previously been linked to nucleotide synthesis^34^, our model does not predict any advantage of such a behavior in the TME.

**Fig. 2:**
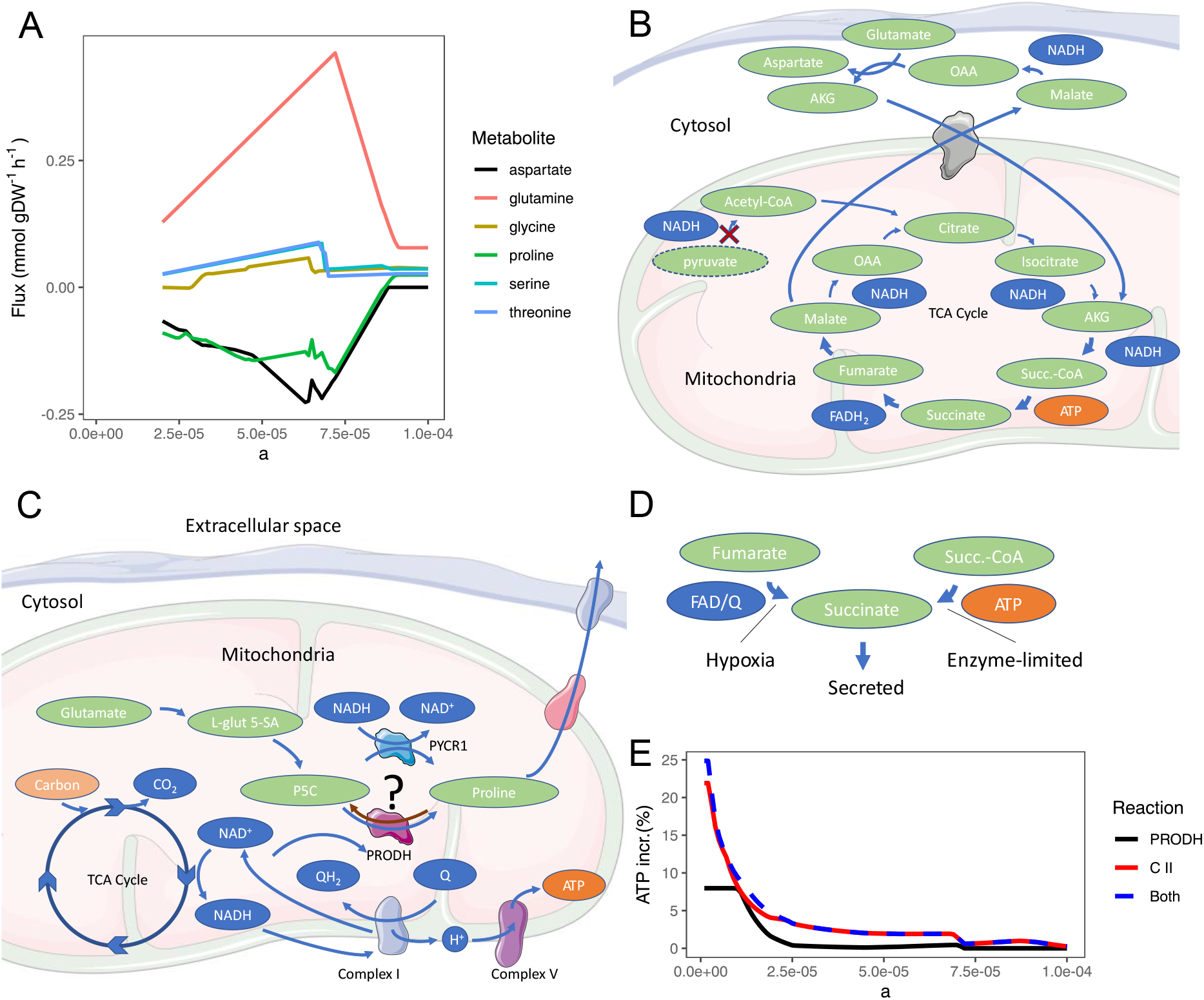
Predicted amino acid metabolism in simulation of cancer cell growth. A. amino acid fluxes from the simulation used in Fig. 1B. Only amino acids with high predicted fluxes are shown. Positive fluxes correspond to cellular uptake and negative to export. B. Predicted “glutamine addiction” mechanism. When OXPHOS is limited by its high enzyme usage, the TCA cycle flux is limited because there is no way for the cell to oxidize the produced NADH/FADH_2_. Any processes that can either limit the production of NADH/FADH_2_ while running the TCA cycle or increase oxidation of NADH/FADH_2_ will thus lead to a possibility to increase the flux through the TCA cycle, and thereby increase the ATP production. Feeding the TCA cycle from glutamate instead of pyruvate shortcuts the TCA cycle, reducing two less NAD+ molecules to NADH, while also preserving pyruvate that can be converted into lactate to oxidize another NADH molecule. The thickness of the arrows indicates the relative magnitude of flux that passes through the reactions, where thicker arrows represent fluxes of greater magnitude. C. Proline metabolism. Generation of proline can help to increase flux through the TCA cycle as well as bypass complex I and thereby increase ATP production. The model predicts two mechanisms for this purpose: 1) Oxidation of NADH through any of the PYCR enzymes (complex I bypass). 2) By running PRODH in reverse. The latter would increase flux through complex I, since Δ^1^-pyrroline-5-carboxylate (P5C) would replace oxygen as the electron acceptor in the electron transport chain, and could therefore in theory increase ATP production by consuming excess NADH and increasing flux through complex V. The brown arrow represents the proline cycle, where PRODH is run in the thermodynamically favorable direction together with the use of a PYCR enzyme, which forms a cycle that can potentially move electrons from NADH to ubiquinone, thereby bypassing complex I without exporting proline. D. Succinate export. Allowing succinate export yields 2 advantages: 1) Complex II can be run in reverse, which highly resembles the effect from running PRODH in reverse. 2) The TCA cycle can be stopped at succinate, reducing production of NADH and FADH_2_, which enables the cell to run the TCA cycle more rounds and thereby produce more ATP. E. Increase in ATP production from allowing the mechanisms explained in C (PRODH in reverse) and D (complex II in reverse). The model is optimized for ATP production, the NGAM is set to 0, and the metabolite uptake bounds are determined by the diffusion model.

The genome-scale metabolic model can be used to study in detail the most optimal amino acid metabolism in different conditions, where the NADH/FADH_2_ oxidation ratio in the ETC plays an important role. For fast growing cells limited by enzymatic capacity, the ratio should be as low as possible to maximize ATP production (complex I bypass), while the ratio should be as high as possible in oxygen-deprived conditions (Note S6). Cells can utilize amino acids in different ways to manipulate this ratio.

“lutamine addiction” is a well-known cancer trait^8,36–38^, where glutamine, the most highly abundant amino acid in blood, is used instead of glucose-derived pyruvate to feed the TCA cycle. Indeed, the model predicts such behavior. Glutamine is converted to glutamate and enters the TCA cycle, producing 3 less NADH per cycle than if pyruvate is used, resulting in increased ATP production, aspartate production, and reduced reaction oxygen species (ROS) generation (Fig. 2B, Note S6). While aspartate export was not observed in a cell culture experiment of the NCI-60 cell lines^39^, aspartate production from glutamine has been shown to exist in pancreatic cancer cell lines using labeling experiments^40^, and is likely disposed of using alternative pathways (Note S6).

Proline secretion has been observed in cell line experiments^34,39^ and the model predicts two different mechanisms by which proline production from glutamate can help to increase ATP production: NADH oxidation through PYCR1 (or related enzymes), which can also form the “proline cycle”^41^ with PRODH to bypass complex I, or by running PRODH in reverse, enabling the use of complex I without oxygen (Fig. 2C, Fig. S9). The latter has to our knowledge not been reported, and it is unclear if such a reaction is thermodynamically favorable *in vivo*, although it has been shown possible in enzyme assays outside cells^42^. In addition, allowing export of succinate enables 1) the reversal of complex II, which catalyzes a similar reaction and has been shown to run in reverse in mammals (Fig. 2D)^43,44^; and 2) interruption of the TCA cycle, thereby bypassing complex II while running succinyl-CoA synthetase to produce ATP (Fig. 2D). The model predicts these two effects to be even more beneficial than PRODH in reverse under hypoxia, although all effects combined yields the highest benefit (Fig. 2E, Fig. S10).

To investigate the behavior of each amino acid in more detail, we simulated access to a fixed influx of a single metabolite and optimized for maximum ATP production, comparing the use of each amino acid as the sole carbon source to that of using lactate under different conditions (Table 1). A few amino acids could be used to generate ATP without oxygen, such as serine which as part of the serine, one-carbon cycle, glycine synthesis (SOG) pathway has been shown to support ATP production and cancer cell proliferation^45^. However, like lactate most amino acids were not useful for producing ATP in that condition. In hypoxia, and without enzyme constraints, PRODH reversal, and succinate export, some benefits to ATP production can be observed from using amino acids as substrates compared to lactate. Allowing reversal of PRODH increases the ATP yield from several amino acids dramatically compared to lactate, and a similar effect is observed for allowing export of succinate, although the benefits vary substantially across different amino acids. Under enzyme-limited conditions, both with and without succinate export, glutamine and glutamate are the carbon sources giving the highest ATP yield, consistent with the strategy depicted in Fig. 2B and the glutamine addiction results reported previously^8^. While such an effect was not observed for hypoxic conditions with low enzyme usage, complex I was bypassed and under enzyme-limited conditions the TCA cycle flux was several times higher for glutamine, yielding a higher total ATP production despite having lower complex V activity (Fig. S11). Furthermore, the PYCR1 reaction (or similar reactions) is used to produce proline, oxidizing 2 equivalents of NADH, which is partly recycled through the proline cycle to reduce ubiquinone, which serves as a complex I bypass mechanism. A previous study showed that glutamine is used to feed the TCA cycle both in hypoxic and normoxic conditions^33^, which is consistent with our results assuming either PRODH in reverse or succinate export in hypoxia.

**Table 1:**
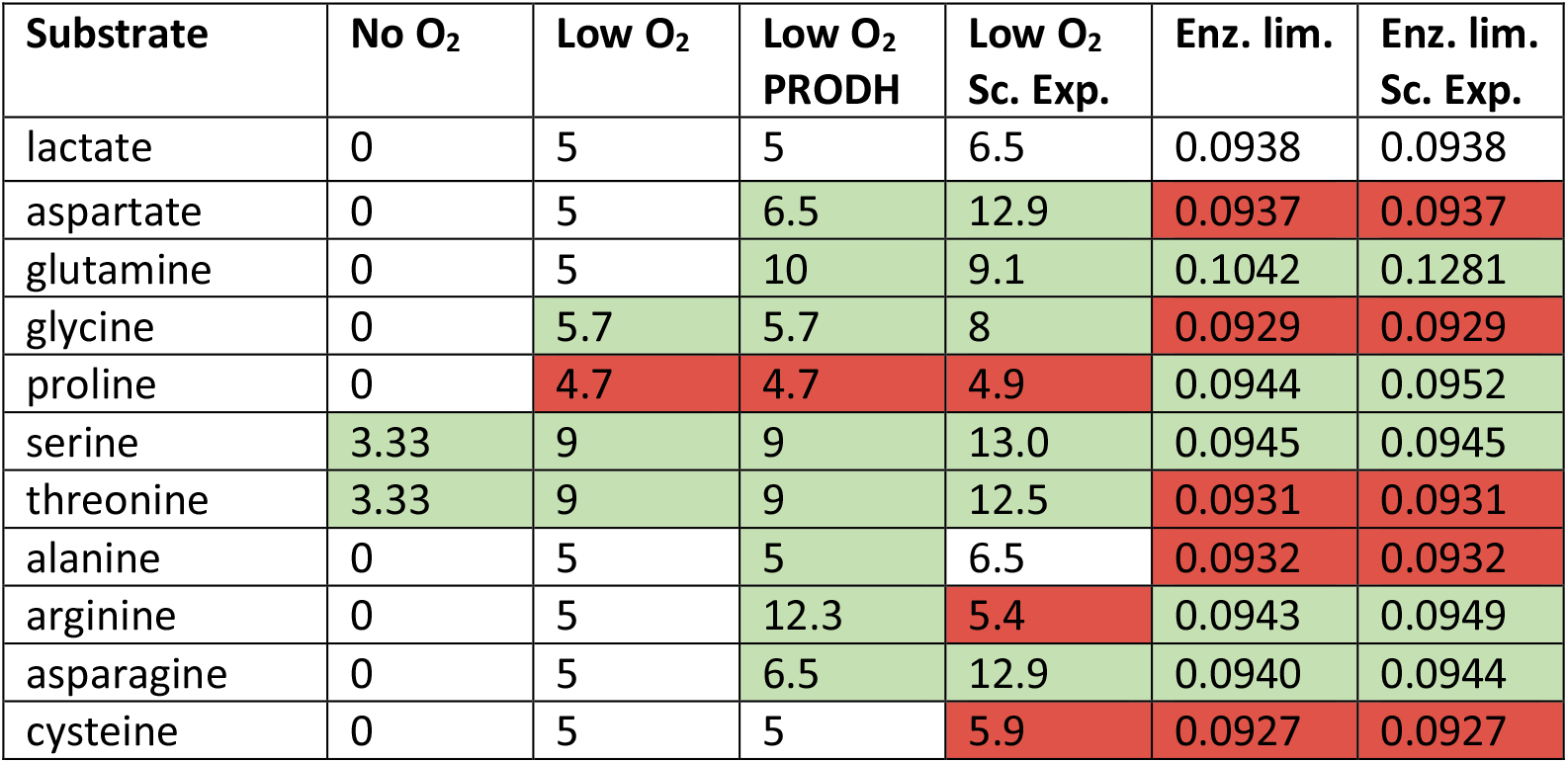
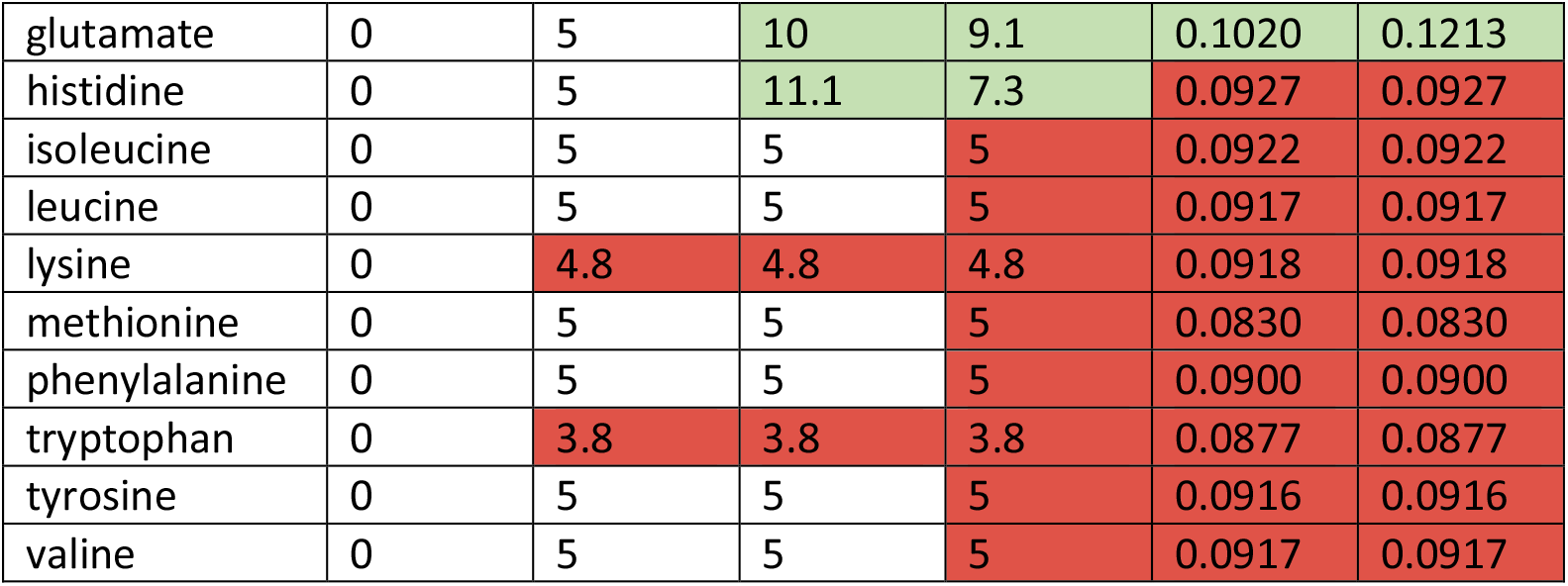
ATP production from amino acids compared to lactate in different settings. The table shows the maximum ATP production (mmol*gDW^−1^h^−1^) given a maximal uptake of 10 mmol*gDW^−1^h^−1^ of a single substrate and varying oxygen availability. Columns: “No O_2_”: No oxygen uptake is allowed, negligible effects from enzyme constraints. “Low O_2_”: Oxygen uptake constrained to 1 mmol*gDW^−1^h^−1^ (which is not enough to fully oxidize any of the substrates), negligible effects from enzyme constraints. PRODH in reverse and succinate export are both blocked. “Low O_2_ PRODH”: Same as “Low O_2_”, but with PRODH in reverse enabled. “Low O_2_ Sc. Exp.”: Same as “Low O_2_”, but with succinate export enabled. “Enz. Lim.”: The total available enzyme pool is constrained to a low value (0.001 g/gDW). Succinate export is blocked. In practice, this also means that the reverse PRODH reaction will not be used. “Enz. Lim. Sc. Exp.”: Same as “Enz. Lim.” but allowing for succinate export. Green background corresponds to a higher ATP production compared to lactate, white to identical, and red to a lower flux. While comparison to using pyruvate as fuel may at first seem more relevant since pyruvate is the alternative fuel for the TCA cycle coming from glycolysis, the pyruvate will be exported as lactate if not used, making comparison to lactate fairer from a redox perspective.

### Simulation of cell type collaborations in the TME

A topic that has drawn some attention the last decade is whether non-cancerous cells in the TME assist cancer cells metabolically by providing them with resources that are advantageous for growth. It has for example been proposed that cancer-associated fibroblasts (CAFs) can provide cancer cells with metabolites such as lactate and pyruvate^10,46^, and it could also be beneficial if tumor-associated macrophages (TAMs) could consume dead cells and cellular debris and produce nutrients for the cancer cells. We sought to investigate these collaboration scenarios in more detail using our diffusion modeling approach. We again constrained metabolite uptake from blood but built a more complex metabolic model consisting of three cell types: cancer cells, fibroblasts, and other cells, where the latter represent cells that are not expected to provide resources to the cancer cells, for example immune cells (Fig. 3A). The exchange of metabolites between the cell types was controlled by providing separate compartments for the interstitium around the fibroblasts and other cells (Fig. 3B).

**Fig. 3:**
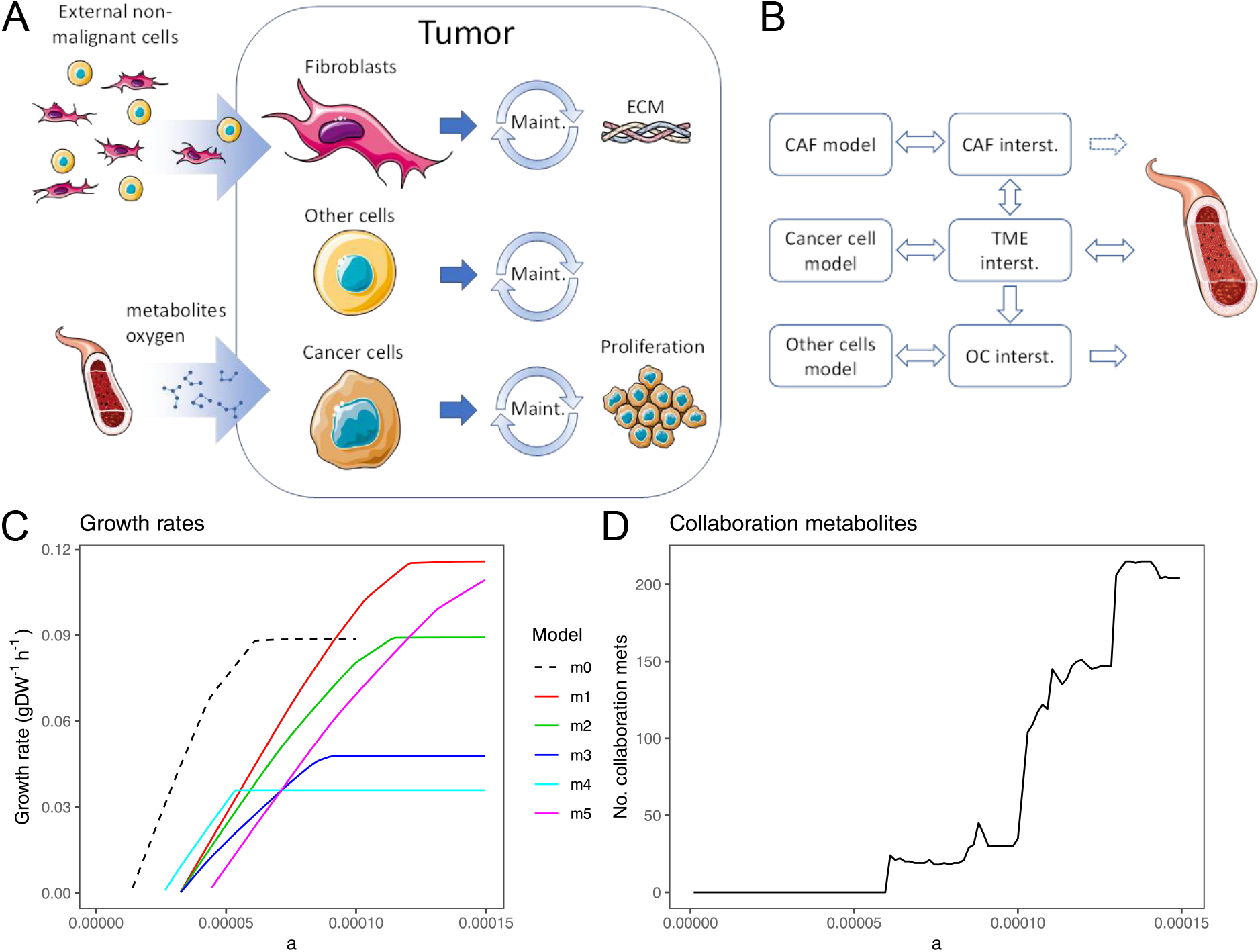
Collaboration between cell types in the TME. A. Modeling setup of the tumor microenvironment. The model contains three cell types: Fibroblasts (CAFs), cancer cells, and other cells (e.g., different types of immune cells). Non-malignant cells are imported to the tumor, resulting in a total biomass reaction consisting of cancer cell growth and extracellular matrix construction. All cell types have the same metabolic network (the curated full Human1 model) with the same NGAM value as used previously, but the fibroblasts were extended with reactions for building the extracellular matrix. B. Compartment setup to enable limitation of cell type collaboration to certain scenarios. The fibroblasts (CAF) and other cells (OC) have their own model compartments for the tumor interstitium. While metabolites can (depending on the simulation) be transported from the CAF interstitium to the cancer cell interstitium, such transport is not possible from the OC insterstitium. C. Effect of collaboration on the specific growth rate of the tumor. D. Number of collaboration metabolites between fibroblasts and cancer cells as a function of a, measured for the m2 model. We identified in total 233 potential collaboration metabolites across the range of explored a values.

The ECM consists mainly of collagens and glycosaminoglycans (GAGs) and is produced by the fibroblasts. Both the composition of the ECM and the fraction of the total tumor dry weight that the ECM constitutes vary largely across tumors. We assumed an ECM composition of 80% collagen (represented by collagen I) and 20% GAGs (represented by heparan sulfate) to reduce the number of flexible parameters. We varied the total ECM fraction of the tumor dry weight and the fraction of each cell type, since these parameters vary substantially between individual tumors. The different modeling configurations are listed in Table 2.

**Table 2.** Different model configurations used in the simulations. The m0 model contains only tumor cells and is identical to the model used in Fig. 1 and 2. ECM fraction represents the weight fraction of the total objective being maximized that the ECM constitutes.

| Model | Cancer cell frac. | Fibr. cell frac. | Other cells frac. | ECM frac. |
| --- | --- | --- | --- | --- |
| m0 | 1 | 0 | 0 | 0 |
| m1 | 0.6 | 0.2 | 0.2 | 0.01 |
| m2 | 0.6 | 0.2 | 0.2 | 0.25 |
| m3 | 0.6 | 0.2 | 0.2 | 0.5 |
| m4 | 0.75 | 0.05 | 0.2 | 0.25 |
| m5 | 0.45 | 0.35 | 0.2 | 0.25 |
| m6 | 0.9699 | 0.0001 | 0.03 | 0.00001 |

It directly follows from the first law of thermodynamics that CAFs cannot increase the total energy available in the TME unless they actively import or utilize material from outside the TME, which is a case we don’t in estigate here. In cases where the growth is limited by nutrient availability only, there is therefore nothing the CAFs can do to help the cancer cells metabolically, given that the cancer cells behave optimally and thereby express the enzymes needed to utilize the available nutrients. However, as nutrient availability increases and the growth becomes limited by enzymatic capacity, it may be possible for the CAFs to increase growth by providing enzymatic capacity to produce metabolites that are more optimal for cancer cell growth.

We first simulated the specific growth rates of the models with different parameter values for ECM and cell type fractions (*m1*-*m5*) and compared them to that of the *m0* model (Fig. 3C). A high fraction of fibroblasts penalized growth at low values of *a* due to the additional ATP costs from non-growth associated maintenance, while the extra enzymatic capacity supplied by a high fibroblast fraction imparts a growth advantage for high *a* values. We also observed that the cost of building the ECM was not negligible in our simulations. However, the addition of ECM to the model objective function did not substantially alter which metabolites were limiting for growth relative to the *m0* model, despite a high protein content (Fig. S12A).

To investigate the potential collaboration between fibroblasts and cancer cells in more detail, we developed an algorithm to identify potential *collaboration metabolites*, defined as metabolites being exported from fibroblasts and imported into cancer cells, for all values of *a* (Fig. 3D, Table S4). As expected, the number increased with increasing *a*, where the enzymatic capacity becomes increasingly limiting for growth. However, many of these collaboration metabolites, such as ATP, are not physiologically meaningful and have not been reported as being exported from fibroblasts.

When limiting the collaboration metabolites to those reported in literature (lactate, pyruvate, free fatty acids, glutamine, ketone bodies, and alanine ^46,47^), and comparing with the model where collaboration is blocked, the growth difference was negligible (Fig. S12B). Although co-injection of tumor cells with fibroblasts in mice can lead to an increased specific growth rate ^48^, we reason that this effect is likely caused by signaling effects or potentially because the fibroblasts complemented deficiencies in the metabolic functionality of these cancer cells. Likewise, mouse experiments have suggested a metabolic collaboration between oxygenated and hypoxic cancer cells^49,50^. We however argue that there is an alternative explanation for these results (Fig. S13).

Although the “other cells” cannot provide the cancer cells with resources in the simulation, they can still minimize their negative impact on growth, which we have termed passive collaboration. We simulated this behavior by maximizing tumor growth using the *m6* model, resulting in ATP generation based on enzyme-demanding processing such as oxidative phosphorylation in the other cells (Fig. S12C). The question is whether the other cells, representing for example immune cells, are helpful for cancer growth, or if the relationship is more competitive. TAMs have however been reported to switch to oxidative phosphorylation when exposed to lactate, which could be seen as a sign of such passive collaboration^51^.

TAMs are reported to have many metabolic roles in the TME, including both passive and active collaboration^51,52^, which would give them the same role as other cells or fibroblasts in our simulation. Macrophages however also have an additional function where they can scavenge dead cells and convert them into metabolites useful to other cells. To investigate to what extent this function can support tumor growth, we used the *m0* model and assumed that 10% of the tumor cells produced from growth die. We assumed that the maximum amount of additional nutrients provided is thus the metabolites that constitute 10% of the produced biomass (Methods), excluding the ATP cost consumed during growth (Table S5). There is a clear growth advantage, which is larger for small *a* values, but the overall effect is relatively small even though no maintenance or metabolite conversion cost is included for the macrophages (Fig. S12D). The reason for the limited utility of such substrates is mainly that in nutrient-deprived conditions all oxygen is already being consumed, and oxygen is required to generate ATP from most scavenged resources. A similar theory to that of scavenging dead cells by macrophages concerns catabolism of organelles and apoptosis in CAFs to provide additional nutrients^10^, where CAFs could then for example be recruited to the tumor. Such a case could provide a small growth advantage similar to the macrophage case but also suffers from the same problem of oxygen limitation and is likely of limited importance.

We also performed these simulations with the blood flow model. In contrast to the diffusion model, a small (negligible) collaboration benefit was observed from collaboration using literature metabolites (Fig. S14A), and the macrophage collaboration utilizing dead cells was more beneficial in this case (Fig. S14B). However, the benefits were still small, confirming our previous conclusion that these metabolic collaboration scenarios are not important for optimizing cancer growth.

## Discussion

During the last decade, metabolic collaboration between cell types in the tumor microenvironment has been thought to increase tumor growth^10–13^, although experimental evidence is largely lacking due to difficulties in measuring and controlling fluxes into and between cell types in an experimental setup. As an alternative, we used genome-scale modeling of the human intracellular metabolic network and integrated measurements of metabolite concentrations in blood and metabolite diffusion coefficients to estimate the maximal increase in specific growth rate from such collaborations at different levels of hypoxia.

The major insight gained from this study was that the metabolic collaboration scenarios between cell types proposed in literature^10–13^ have a negligible effect on tumor growth. Although fibroblasts have been shown to export lactate^53^ and some tumors have been shown to take up lacate^54^, no direct evidence is presented that such a behavior is beneficial for growth. In addition, scavenging of dead cells by macrophages or other forms of cell catabolism such as catabolic behavior of fibroblasts ^13^ only yield a marginal benefit to growth due to a shortage of oxygen. However, the signaling interplay between stromal cells and cancer cells is most likely important for changing the metabolism of tumor cells^13^.

An additional insight gained from this study is that ATP production is most likely the limiting factor for growth for cancer cells in the tumor microenvironment, and that the ATP production is limited by either lack of nutrients and oxygen, by limitations in enzymatic capacity, or a combination of both. Other factors not considered in our model may also play a role in limiting growth, such as extensive ROS production or low pH in the interstitial fluid. The high need for ATP is implicitly supported by literature; most of the carbon from glucose and glutamine imported by cells is not incorporated into the biomass^55^, suggesting that it is used for ATP production.

The third important insight from this study concerns the use of amino acids for energy production and the varying ways in which the TCA cycle can be used. Our model for example predicts glutamine addiction^8,56,57^ and gives plausible explanations to why glutamine is a more optimal substrate than pyruvate for the TCA cycle at different levels of hypoxia. We also predict potential benefits from running PRODH in reverse in hypoxia, although it is unclear if such a reaction is thermodynamically favorable, and predict a potential 20% increase in ATP production from running mitochondrial complex II in reverse, which has been shown to take place both in cell lines and in vivo^44^.

Altogether, our results suggest that there is no benefit for tumors to develop a metabolic collaboration between cell types. Such metabolic collaborations may still exist, where a cell type can complement deficiencies in tumor cells, which has for example been observed during glutamine deprivation^58^. However, such a configuration does not improve growth beyond what enabling the needed pathway directly in the tumor cells would yield. Our modeling approach allows for investigation of the optimal behavior for cells in the TME under the assumptions that the diffusion and blood flow models are based upon. While these models are approximations of the true influx of metabolites into tumors, they can still serve well for future investigations of different metabolic phenomena in the tumor microenvironment, and with a systems biology approach unravel questions that are difficult to address using experiments.

## Methods

### Model preparation

We used the genome-scale metabolic model Human1 (version 1.12) with added enzyme constraints (using Gecko Light, see Note S1). The model was manually curated by inactivating 25 reactions, since they led to unrealistic fluxes (Table S6). We removed all reactions related to amino acid triplets and drug metabolism. Furthermore, we removed all reactions that could not carry flux given the metabolites available in blood and am’s media, which were identified using a modified version of the function “ha e lu “in A oolbo, except for the macrophage simulation where these reactions were retained since new metabolites were added as input. Furthermore, a non-growth associated maintenance (NGAM) value of 1.833 mmol*gDW^−1^h^−1^ was collected from literature ^20,21^ and added to the model as a lower bound on the reaction MAR03964, representing hydrolysis of ATP to ADP.

### Combined model

The combined model contained three cell types: CAFs, cancer cells, and other cells, where the latter refers to cells that do not collaborate with the cancer cells, for example some types of immune cells (Fig. 3A). All three cell types were represented by a separate instance of the curated Human1 model, but the CAFs were extended with reactions for building the extracellular matrix. Each cell type had its own interstitium compartment, where cancer cells had the TME interstitium compartment, which receives all metabolite influx from blood (Fig. 3B). This setup enabled control of metabolite flux between cell types, where the CAFs and cancer cells could exchange metabolites but the flux of metabolites between other cells and cancer cells was unidirectional towards the other cells. The tumor biomass reaction, which was used as the FBA objective, contained tumor cell growth and construction of ECM, where the latter was synthesized by the CAFs. CAFs and other cells were assumed to be recruited to the tumor, and the biomass reactions of these cells were therefore not included in the total tumor biomass reaction. The ECM was composed of collagen I and GAGs (represented by heparan sulfate) (Table S2). The protein cost in the ECM biomass reaction also included an ATP cost for polymerizing amino acids into peptides. There was a small enzyme usage cost added for transportation of metabolites from the fibroblast interstitium compartment to the TME interstitium, with the purpose of avoiding unnecessary transportation of metabolites between cell types (which could add additional collaboration metabolites).

### Modeling amino acid metabolism

We used a special modeling configuration to generate the results in Table 1. For all setups, we first blocked the input of all metabolites and removed the protein usage constraint. We then set the upper bound of the input exchange reactions of water, phosphate, oxygen, and H^+^ to infinity. The objective function was then set to maximize the flux through the ATP hydrolysis reaction (MAR03964), and the NGAM ATP cost was removed. For each metabolite, we then set the upper bound of the input exchange reaction of that metabolite to 10 mmol*gDW^−1^h^−1^. The optimization was run in several settings: ***“****Low O*_*2*_*”:* Here, the upper bound of oxygen was set to 1 mmol*gDW^−1^h^−1^. *“Low O*_*2*_, *no PRODH”:* Same as *“Low O*_*2*_*”* but the re erse reaction (“A 03838”) is also blocked. *“No O*_*2*_, *no PRODH”:* Same as *“Low O*_*2*_, *no PRODH”*, but the oxygen upper bound is set to zero. *“Enzyme Lim.”:* The protein usage was constrained to 0.001 g/gDW.

### Flux Balance Analysis (FBA)

Flux balance analysis was performed using RAVEN Toolbox v. 2.7.5^59^. We used Gurobi v. 9.5.0 as the solver and set the numerical tolerances to 10^−9^.

### Flux Variability Analysis (FVA) to determine growth-limiting metabolites

To identify growth-limiting metabolites, we first optimized the model for growth (maximization of flux through the biomass reaction). The lower bound of the biomass reaction was then set to the maximum specific growth rate (subtracted by 10^−4^ to avoid numerical issues in the solver), followed by an optimization for each metabolite *i* of interest, where the flu of the metabolite’s import exchange reaction was minimized, resulting in the minimum uptake rate *u*_*m,i*_ needed to sustain the maximum specific growth rate. The minimum required fraction of the metabolite, *f*, was then calculated as

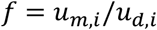

where *u*_*d,i*_ is the maximum uptake rate of the metabolite as defined by the diffusion model.

### Identification of collaboration metabolites

Potential collaboration metabolites for each value of *a* were identified using an iterative algorithm on the m2 model. The algorithm in each iteration finds the potential collaboration metabolites, defined as metabolites that are exported from the fibroblasts and imported into the cancer cells when optimizing the model for total tumor growth using FBA. These metabolites are added to a list of potential collaboration metabolites and are then blocked from further collaboration by zeroing the upper bound of the transport reaction from the fibroblast interstitium compartment (f_s) to the TME interstitium compartment (s). The procedure is then repeated to identify new potential collaboration metabolites to add to the list, until no more such metabolites are found. Metabolites with infinite availability, such as water and H^+^, are excluded. We don’t in estigate if potential collaboration metabolites can actually contribute to growth since such an effect may be complex and is not easily identified.

### Macrophage collaboration

To estimate the increase in uptake bound for metabolites from macrophage collaboration, we first derived the metabolite composition of the cells from the reaction “biomass_human”, ignoring A (since it is spent during growth) and some lowly abundant metabolites. A few metabolites that did not have exchange reactions were either changed to a similar compound or removed. We then for each value of *a* first optimized for growth. The additional metabolite availability *u*_*add,i,x*_ for metabolite *i* at *a*=*x* was then calculated as

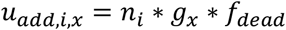

where *n*_*i*_ is the molar composition of metabolite *i* in the biomass reaction (mmol/gDW), *g*_*x*_ is the specific growth rate at *a*=*x*, and *f*_*dead*_ is the fraction of cells produced that die, assumed to be 0.1 in the simulations. he fatty acid pool was con erted to the metabolites “cholesterol” and “A blood pool in” he metabolites and their concentrations are a ailable in able

### Software

The data was analyzed using MATLAB R2019b and R version 3.6.1. MATLAB was used for all analyses and the figures were generated in R. To ensure the quality of our analyses, we verified and validated the code using a combination of test cases, reasoning around expected outcome of a function, and code review. The details of this activity are available in the verification matrix available with the code.

## Declarations

### Availability of data and materials

The model Human1 is available in GitHub (https://github.com/SysBioChalmers/Human-GEM). *k*_*cat*_ values are indirectly available in the Gecko Light repository in GitHub (https://github.com/SysBioChalmers/GeckoLight). Metabolite concentrations and diffusion coefficients are available in Table S3. The processed data and source code are available in Zenodo: https://doi.org/10.5281/zenodo.7358685. The source code is also available in GitHub: https://github.com/SysBioChalmers/TMEModeling.

### Authors’ contribution

J.G, J.R, and J.N. planned the project. J.G. and A.H. developed the diffusion model. J.G., F.R., D.B., S.P., and O.D. investigated and drew conclusions about the amino acid metabolism. J.G. wrote all software and made all analyses and figures. J.G. wrote the draft manuscript. All authors reviewed and edited the manuscript. J.R. and J.N. supervised the project. J.N. acquired funding for the project.

## Acknowledgements

We thank Stig Larsson for reviewing the diffusion model and Gunjan Purohit for help with investigating amino acid metabolism.

