## Supplementary information for "Metabolic collaboration between cells in the tumor microenvironment has a negligible effect on tumor growth"

### Supplementary Figures

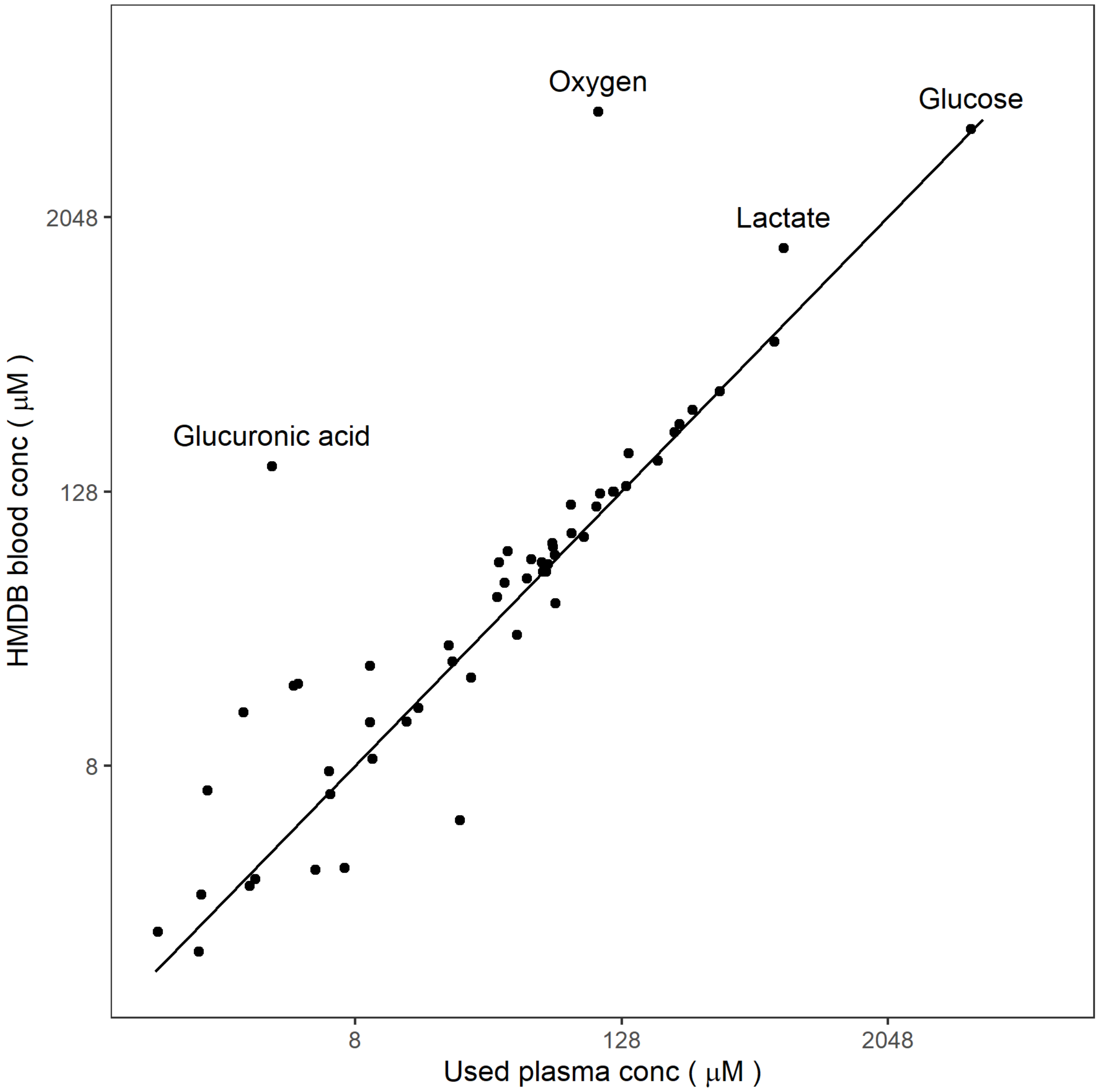


***Fig S1: Validation of used blood plasma metabolite concentrations.*** *The figure shows a comparison between HMDB[1,2] and the data from Akinci et al (for glucose, HMDB value used in simulations), Siggaard-Andersen et al (for oxygen), and Harada et al[3–5]. The line shows the ideal relationship between the data sources, it is not a linear fit to the points. Only metabolites with data from both sources where the metabolite name could be mapped to the model are shown.*

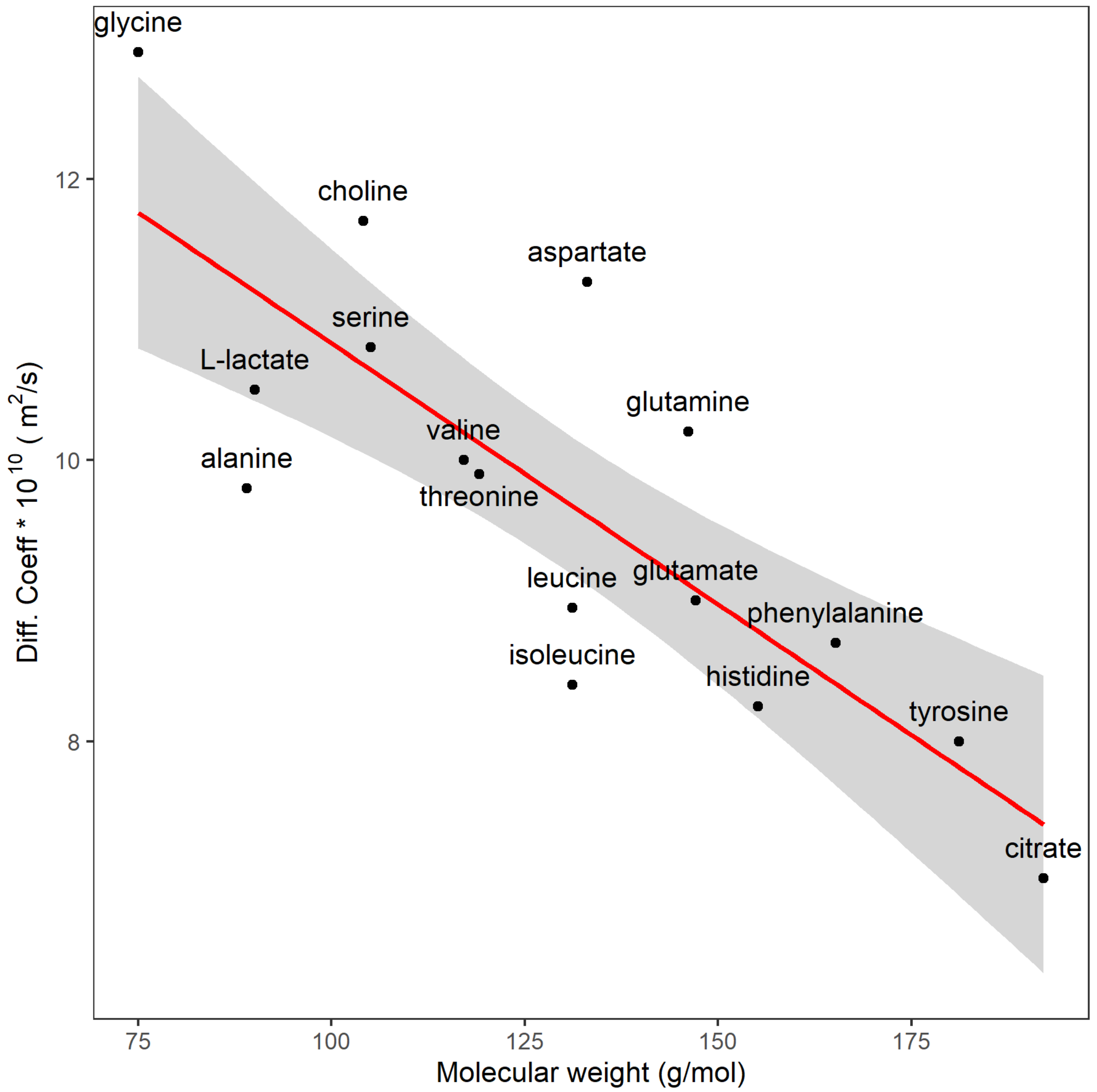


***Fig. S2: Linear model for estimation of diffusion coefficients from molecular mass.*** *The model is used to estimate the diffusion coefficients for metabolites where no such value is available. Oxygen and albumin are not included in the model since a linear model is only valid within a limited molecular mass range.*

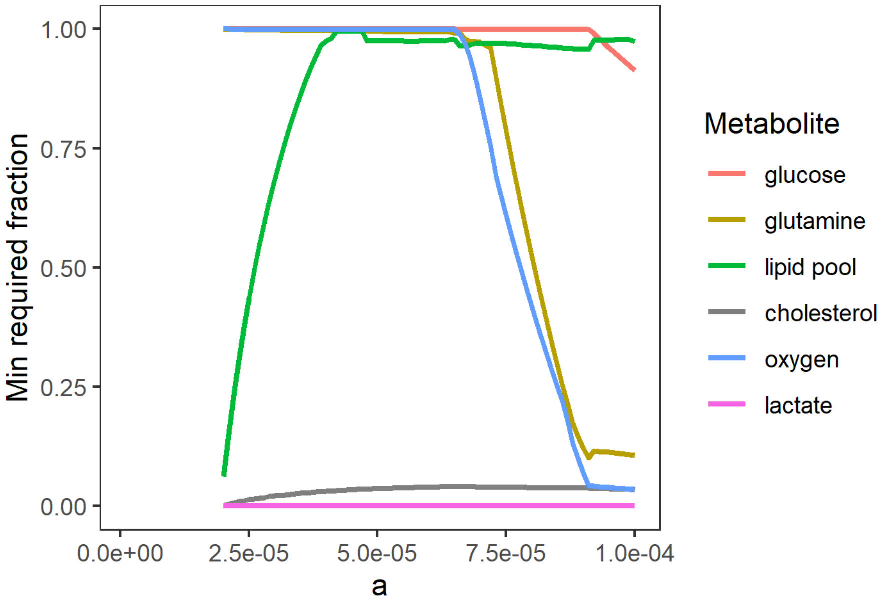


***Fig. S3: Growth limitation of metabolites when simulating tumor cell growth.*** *Investigation of which metabolites are limiting for growth at different values of* a*. The simulation is performed by fixing the specific growth rate at the maximum possible specific growth rate, followed by a minimization of the uptake of a specific metabolite (FVA). The "min required fraction" represents the minimum required uptake rate of the metabolite divided by the maximal possible value – a value of 1 thus represents that a metabolite is limiting for growth.*

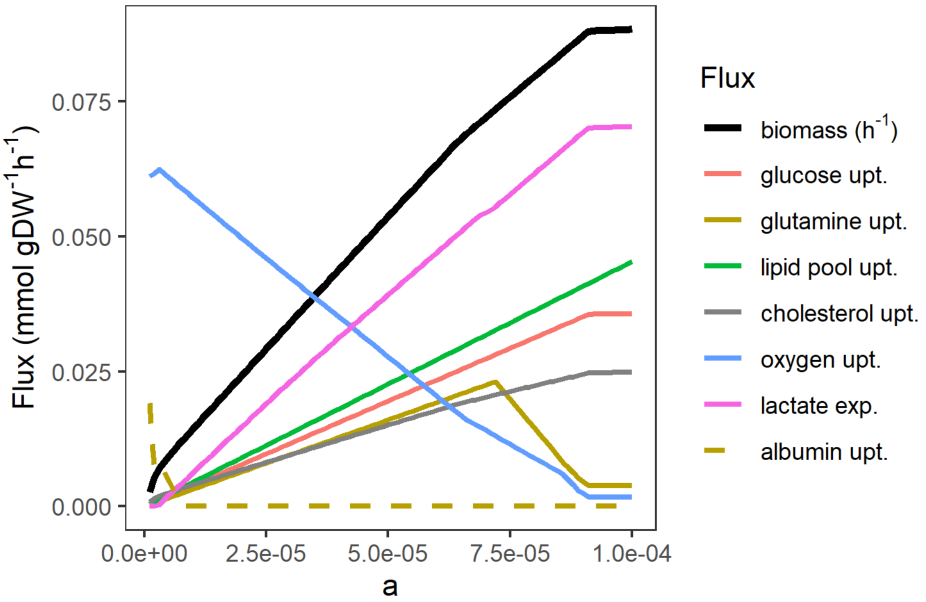


***Fig. S4: Simulation of tumor cell growth without oxygen constraints.*** *Identical to the simulation in Fig. 1B in the main text except that oxygen uptake is unconstrained. Albumin uptake is also added to the figure. The albumin uptake rate is multiplied by 200 to fit the scale of the other fluxes, while other metabolites are scaled the same way as in Fig. 1B in the main text. Albumin was useful for growth at small* a *values since enzyme usage was not limited at such* a *values, allowing for full oxidation of the amino acids for the purpose of ATP generation, which overcomes the ATP cost associated with digestion of albumin into amino acids.*

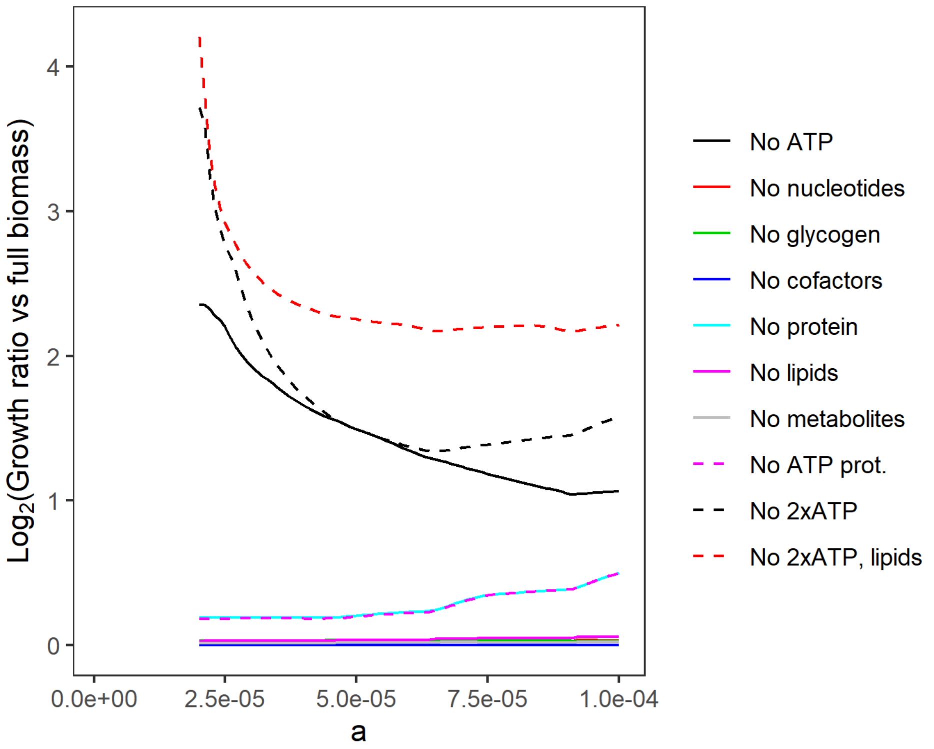


***Fig S5: Change in specific growth rate when removing parts of the biomass reaction.*** *The figure shows the specific growth rate ratio between models with reduced and original biomass reaction. In all cases, the model is optimized for growth. "No ATP prot." refers to removal of the ATP cost from turning amino acids into proteins, while "No 2xATP" refers to having both the protein generation ATP cost and the direct ATP cost removed from the biomass reaction. For "No 2xATP, lipids", the consumption of lipids have also been removed in addition to the other two factors. ATP generation is the limiting factor for all values of* a*. With ATP costs removed, the availability of lipids became limiting, while the direct use of amino acids for protein synthesis was small compared to the amounts available.*

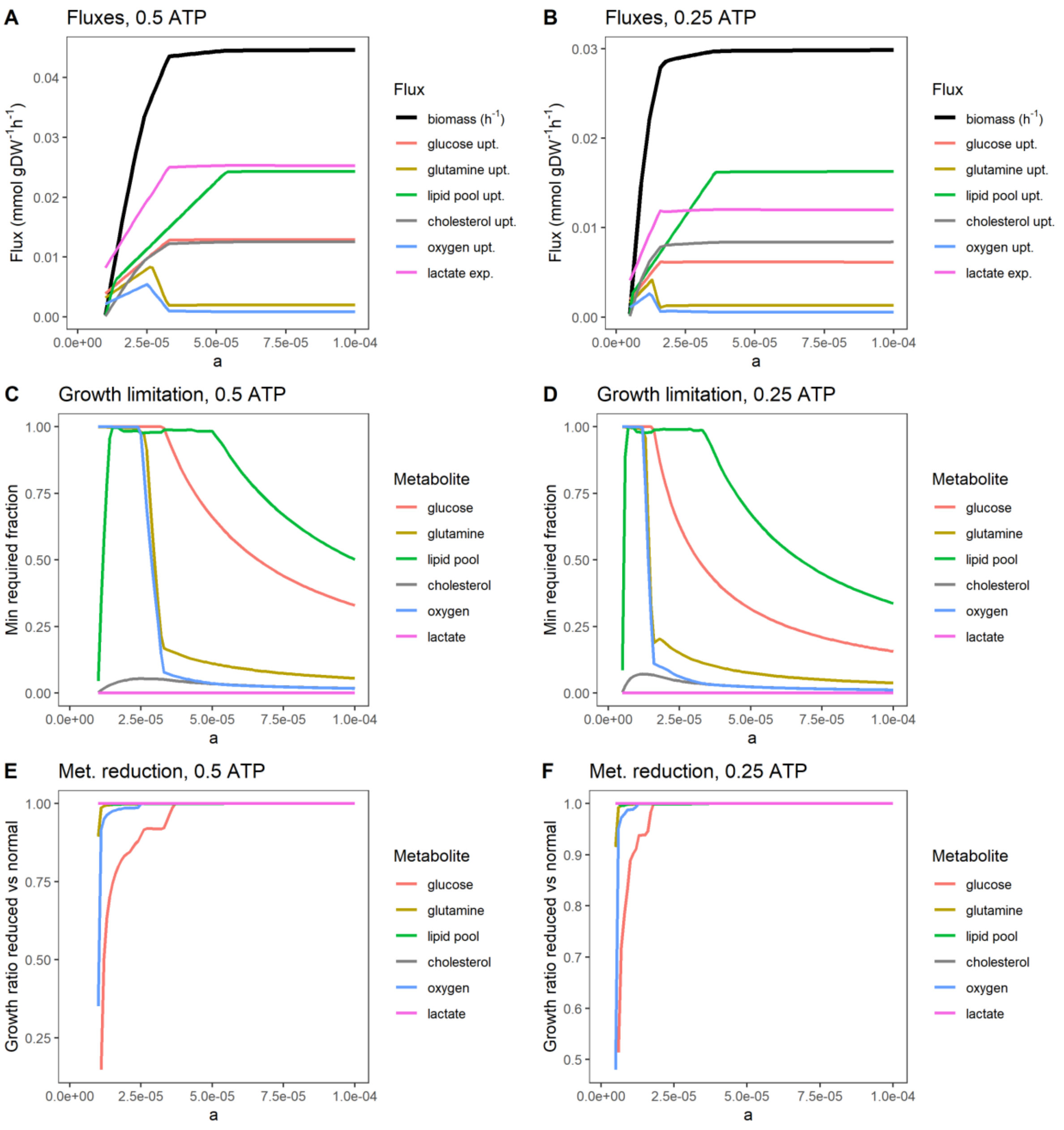


***Fig S6: Metabolites limiting for growth when the direct ATP cost of the biomass function and NGAM are reduced to 50% and 25% of the original values.*** *A-B. Specific growth rate and metabolite uptake/export. C-D. Flux variability analysis showing how much each metabolite can be reduced while retaining growth. E-F. Specific growth rate reduction from reducing the maximum uptake rate of a single metabolite to 90%.*

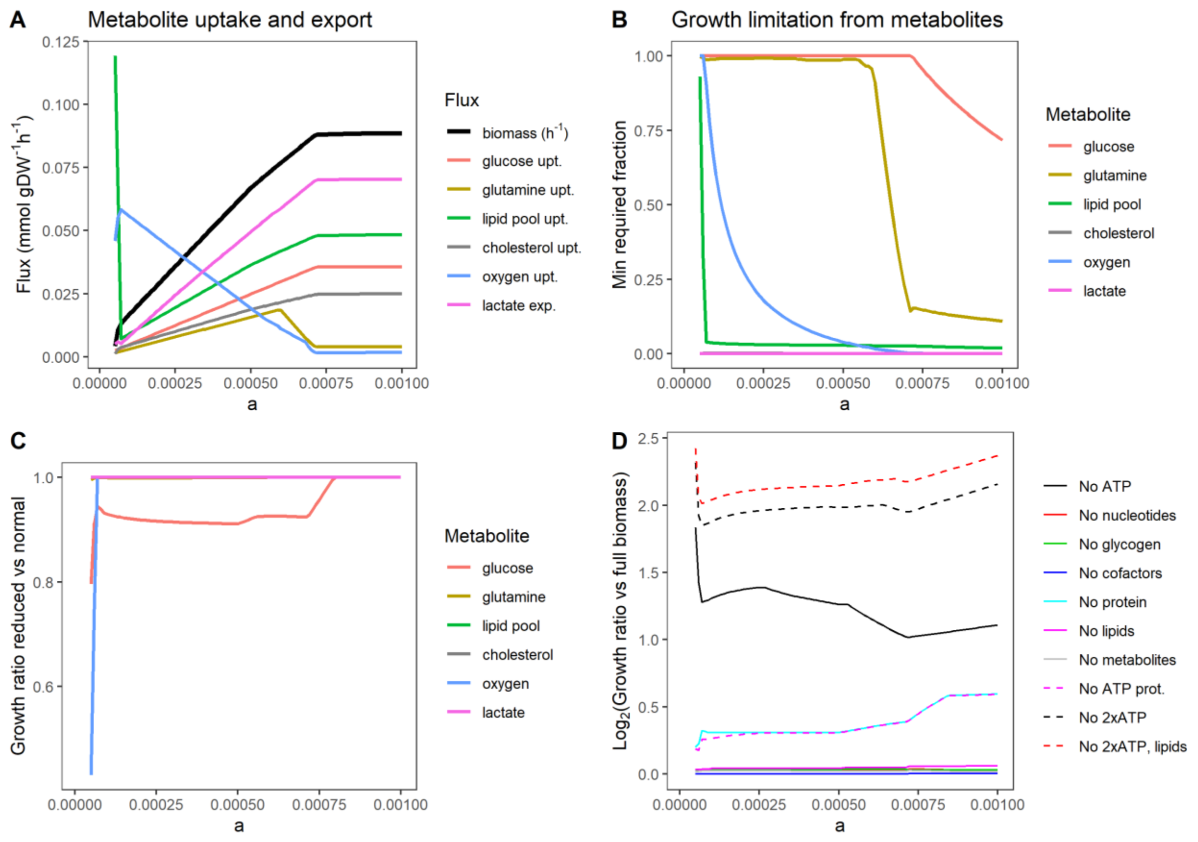


***Fig S7: Metabolic behavior for optimal growth using the blood flow model.*** *A. Specific growth rate and metabolite uptake/export (see Fig. 1B in the main text for comparison with the diffusion model). B. Growth limitation of metabolites when simulating tumor cell growth (see Fig. S3 for comparison with the diffusion model). The simulation is performed by fixing the specific growth rate at the maximum possible specific growth rate, followed by a minimization of the uptake of a specific metabolite (FVA). The "min required fraction" represents the minimum required uptake rate of the metabolite divided by the maximal possible value – a value of 1 thus represents that a metabolite is limiting for growth. C. Specific growth rate reduction from reducing the maximum uptake rate of a single metabolite by 10% (see Fig. S4 for comparison with the diffusion model). The effect from glutamine, lipid pool, cholesterol, and lactate is negligible or non-existent. D. Change in specific growth rate when removing parts of the biomass reaction (see Fig. S6 for comparison with the diffusion model). The figure shows the specific growth rate ratio between models with reduced and original biomass reaction. In all cases, the model is optimized for growth. "No ATP prot." refers to removal of the ATP cost from turning amino acids into proteins, while "No 2xATP" refers to having both the protein generation ATP cost and the direct ATP cost removed from the biomass reaction. For "No 2xATP, lipids", the consumption of lipids have also been removed in addition to the other two factors. ATP generation is the limiting factor for all values of* a. *With ATP costs removed, the availability of lipids became limiting, while the direct use of amino acids for protein synthesis was small compared to the amounts available.*

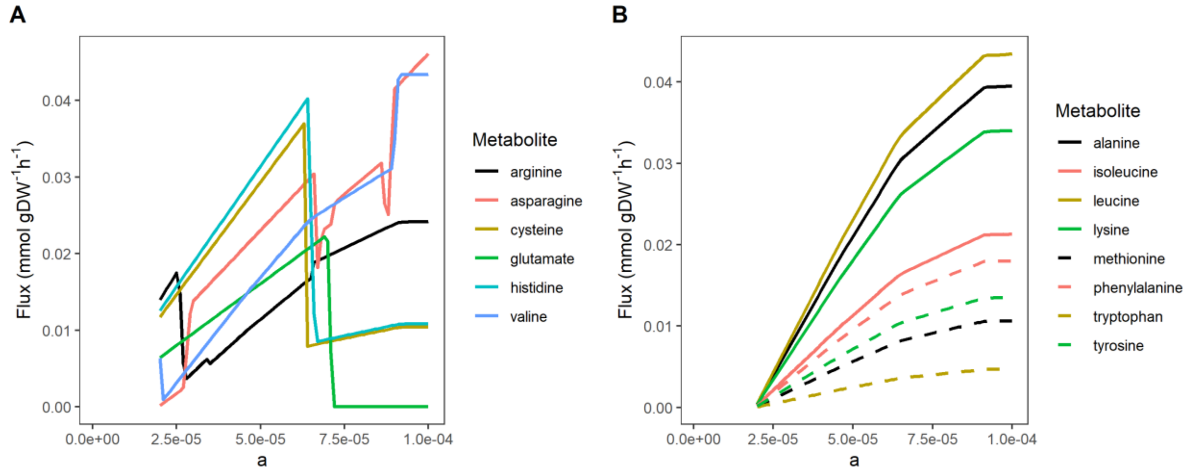


***Fig S8: Uptake of amino acids in the tumor microenvironment simulation.*** *A. Uptake of amino acids exhibiting low but varying uptake in a simulation of tumor cell growth with metabolite uptake constraints derived from the diffusion model. B. Uptake of amino acids that are mainly used for protein production in a simulation of tumor cell growth with metabolite uptake constraints derived from the diffusion model. These figures are similar to Fig. 2A in the main text.*

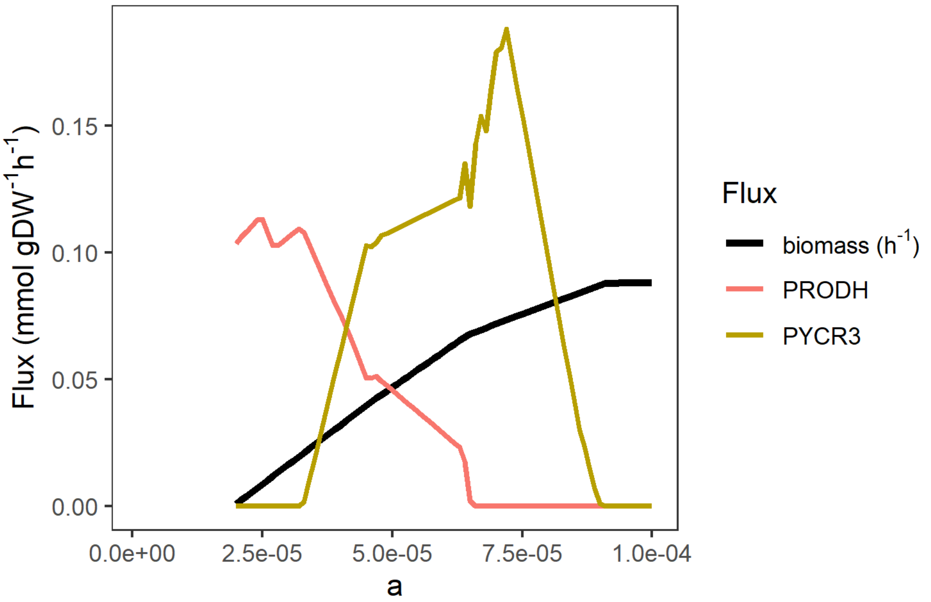


***Fig S9: Fluxes through two proline reaction pathways used by the model for increasing growth.*** *The PRODH curve represents the reverse flux through PRODH. We note that the model chooses to use PYCR3 here, while we in Fig. 2C in the main text has depicted the flux through PYCR1. The choice of PYCR enzyme by the algorithm is largely arbitrary; any of the PYCR enzymes can be used, although PYCR3 activates extra transporters since it resides outside mitochondria.*

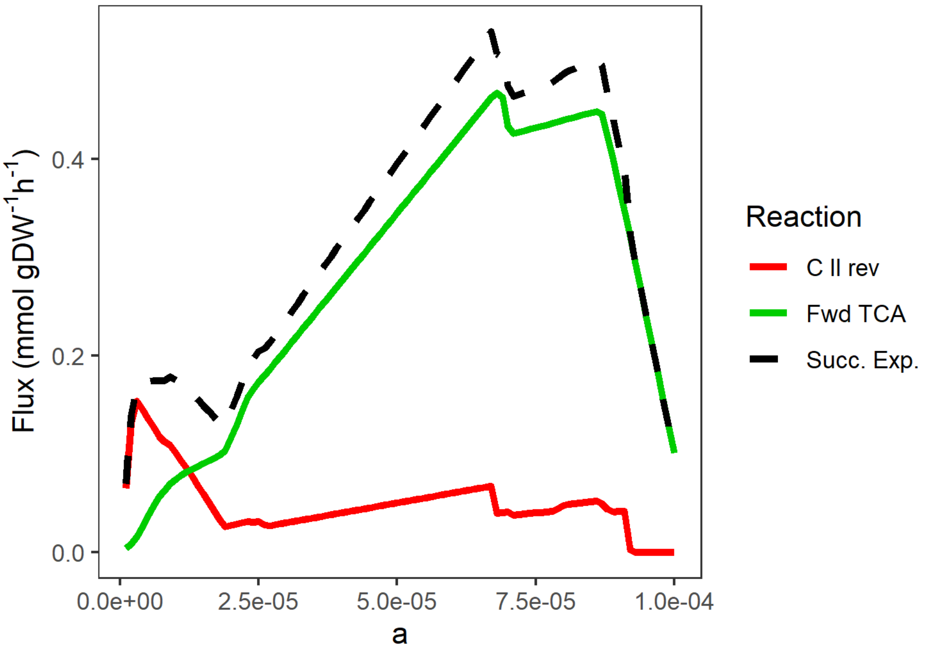


***Fig S10: The source of succinate export.*** *Succinate export and the fluxes of complex II in reverse and succinyl-CoA synthetase in a simulation similar to that of Fig. 1B in the main text but allowing for succinate export.*

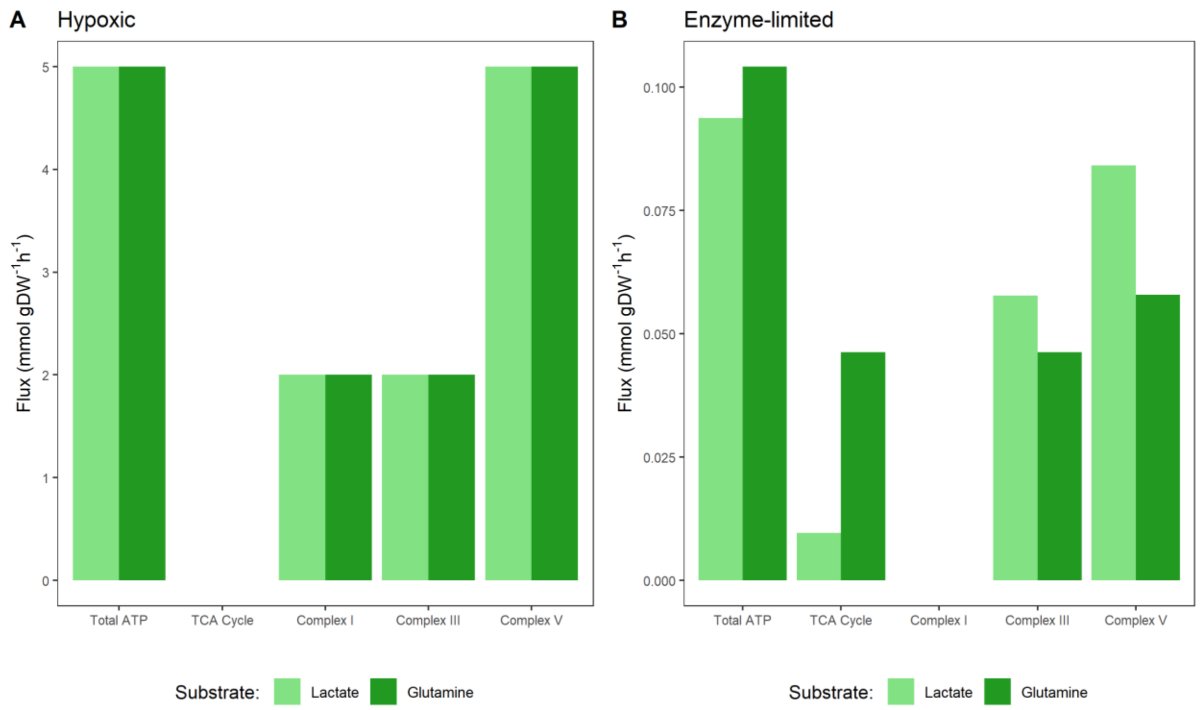


***Fig S11: Differences in ATP production from the substrates lactate and glutamine.*** *A. Hypoxic conditions with the reverse PRODH reaction blocked. B. Enzyme-limited conditions.*

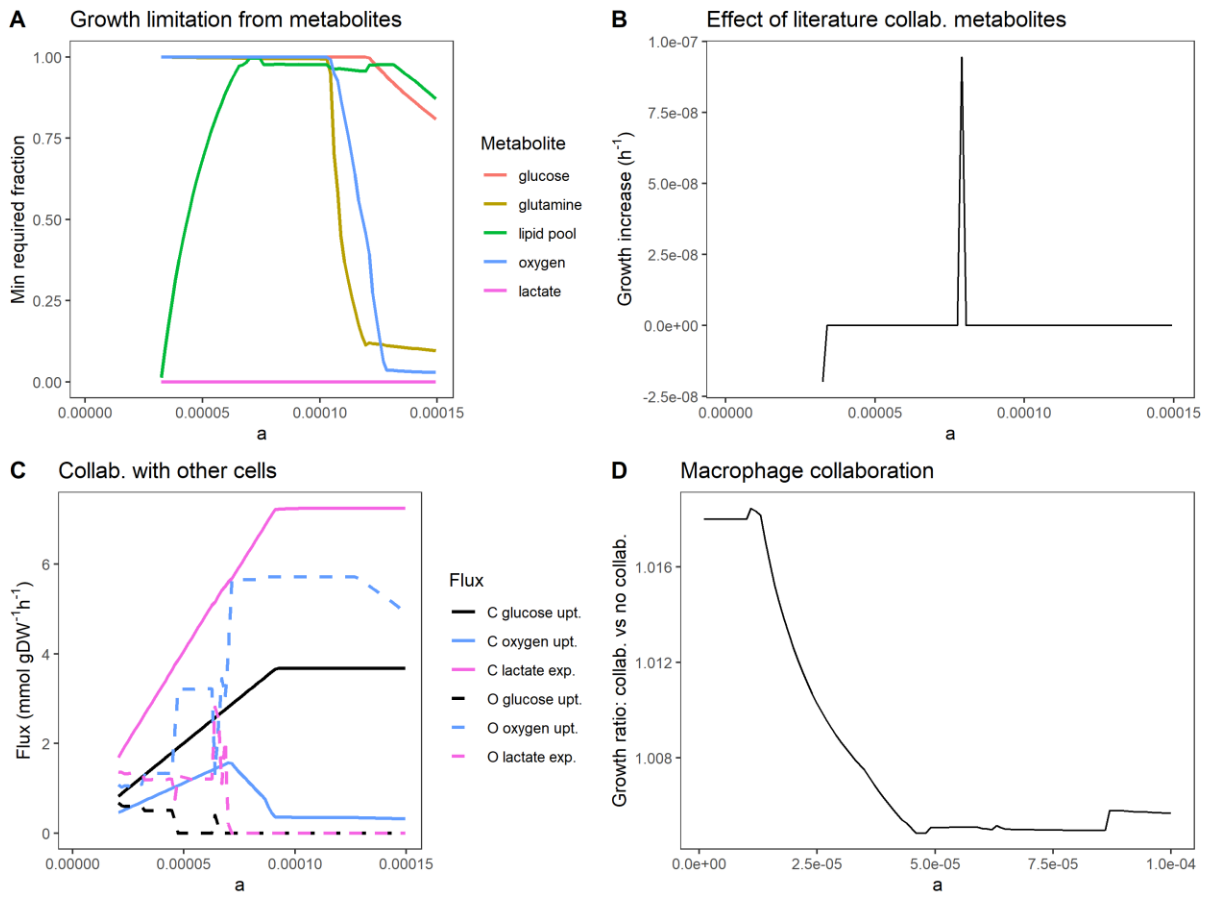


***Fig. S12: Additional plots for the combined model.*** *A. Investigation of which metabolites are limiting for growth at different values of* a *for the m3 model, which has a high ECM content. The simulation is performed by fixing the specific growth rate at the maximum possible specific growth rate, followed by a minimization of the uptake of a specific metabolite. The min required fraction represents the minimum required uptake rate of the metabolite divided by the maximal possible value – a value close to 1 thus represents that a metabolite is limiting for growth. B. Maximum increase in specific growth rate by collaboration with CAFs allowing only metabolites previously identified in literature to be sent from the CAFs to the tumor cells. C. Metabolite uptake rates in a passive collaboration between cancer cells and other cells. The m6 model is used, which has negligible fractions of both fibroblasts and ECM, and only a small fraction of other cells, which due to the low availability of oxygen is beneficial for showing shifts in oxygen use between cell types for different values of* a*. The simulation shows the expected behavior; the other cells consume little glucose while instead consuming the metabolites that are least useful for the cancer cells. These are a large pool of compounds such as citrate and linoleate together with oxygen and are used to generate the maintenance ATP needed to sustain the cells. D. Maximum increase in specific growth rate by collaboration with macrophages that reuse 10% of the produced biomass (dead cells).*

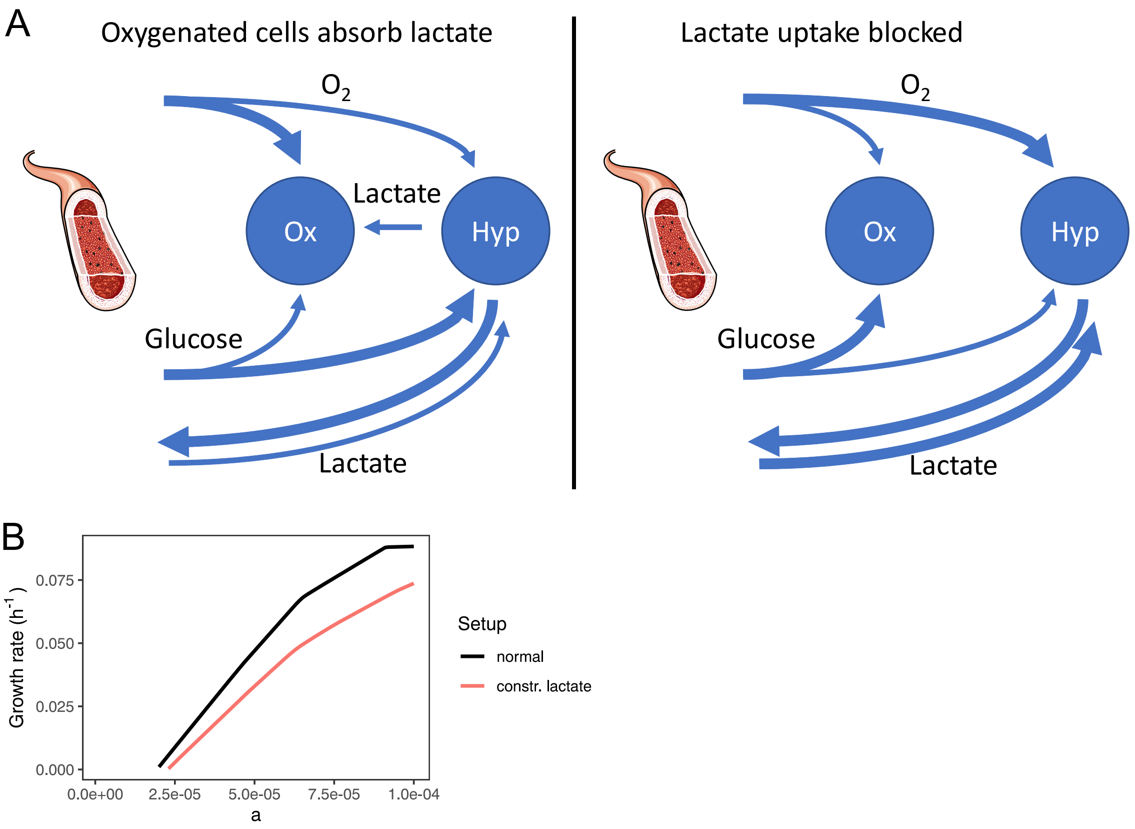


***Fig. S13: Alternative explanation to collaboration between oxygenated and hypoxic cancer cells.*** *A. It has been reported that oxygenated cancer cells (from well oxygenated regions of the tumor, representing a large value of* a*) collaborate with hypoxic cancer cells (from hypoxic regions, representing a lower* a *value) by consuming lactate*[6,7]*. Less glucose will then be spent in the well oxygenated regions and can diffuse to the hypoxic areas, and the necrotic regions in mouse tumor was shown to increase when lactate uptake was blocked*[6]*. A problem with this theory is that a corresponding increase in oxygen uptake is expected in the well oxygenated regions, since more oxygen is required to produce the same amount of ATP from lactate as compared to glucose, and less oxygen therefore will diffuse to the hypoxic regions, potentially counteracting the beneficial effect. An alternative theory could be that the increase in necrotic regions is related to the experimental setup where the lactate uptake was blocked by an inhibitor. The lactate uptake is then reduced in the whole body, likely resulting in higher lactate levels in blood, and therefore an increase in lactate influx to the tumor and consequently a decrease in pH in the extracellular compartment. The internal pH in cells must stay above a certain level for cell survival*[7–10]*, and it has been proposed that lowering the external pH makes active regulation of internal pH increasingly demanding, leading to a higher ATP demand for the cells*[8,11]*, and consequently, an increase in the necrotic regions. In addition, it has been shown that mammalian cells that are not under stress generally tend to take up lactate whenever available*[12]*, for example to regulate the pH of blood and tissue, and cancer cells may have simply retained this behavior from healthy cells. We conclude that although these results at first appear contradictory to our results, the increase in necrotic regions could just as well be explained by increased pH caused by the inhibitor and not due to interrupted collaboration between these cancer cells. B. Investigation of the impact on growth from constraining lactate output (and thereby production) for the purpose of increasing extracellular pH. The maximum lactate output was constrained to half of the glucose uptake bound, which showed a large negative impact on growth and a small increase of the necrotic region, further motivating that low extracellular pH can affect growth and survival negatively.*

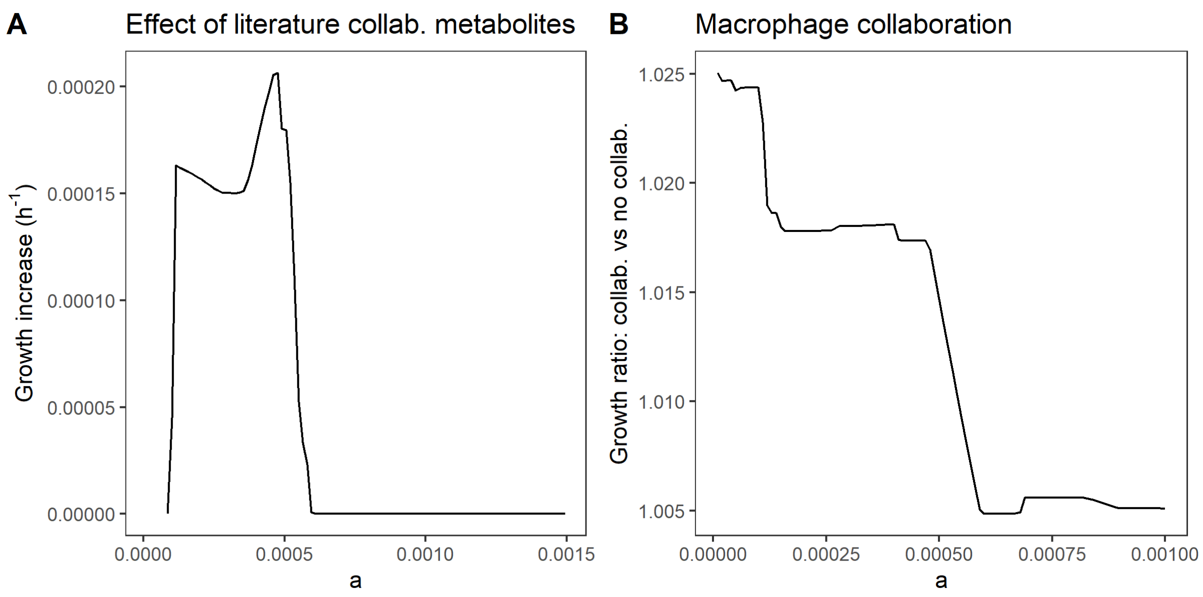


***Fig. S14: Results from the combined model using the blood flow model.*** *A. Maximum increase in specific growth rate by collaboration with CAFs allowing only metabolites previously identified in literature to be sent from the CAFs to the tumor cells. B. Maximum increase in specific growth rate by collaboration with macrophages that reuse 10% of the produced biomass (dead cells).*

### Supplementary Tables

#### Table S1 – Additional blood metabolite candidates from HMDB

Available as a separate Excel sheet.

#### Table S2 – Metabolites from HMDB that could not be mapped to the model

Available as a separate Excel sheet.

#### Table S3 – Consensus metabolite concentrations in blood and corresponding diffusion coefficients

Available as a separate Excel sheet.

#### Table S4 – Potential collaboration metabolites between fibroblasts and tumor cells identified using the m2 model

Available as a separate Excel sheet.

#### Table S5 – Additional metabolite availability from dead cells

Available as a separate Excel sheet.

### Table S6 – Reactions blocked in Human1 for the simulations

The following reactions were blocked in Human1 as a curation step since they gave rise to unrealistic fluxes.

| Blocked reactions | Reaction equation |
|---|---|
| MAR08759 | 2 hypotaurine[c] + O2[c] => 2 taurine[c] |
| MAR02779_REV | glucose[c] + H+[c] + NADH[c] => D-glucitol[c] + NAD+[c] |
| MAR03996 | cystine[c] + H+[c] + NADH[c] <=> 2 cysteine[c] + NAD+[c] |
| MAR06965 | dehydrospermidine[c] + H+[c] + NADH[c] <=> NAD+[c] + spermidine[c] |
| MAR13081[*] | 4 ferrocytochrome C[m] + 7.92 H+[m] + O2[m] => 4 ferricytochrome C[m] + 1.96 H2O[m] + 0.02 O2-[m] + 4 H+[i] |
| MAR08981 | NAD+[x] + malate[x] <=> H+[x] + NADH[x] + OAA[x] |
| MAR03167 | urate[c] => H+[c] + urate radical[c] |
| MAR12019 | leukoaminochrome[c] + O2[c] => H2O2[c] + 2,3-Dihydro-1H-Indole-5,6-Dione[c] |
| MAR01706 | 3alpha,7alpha-dihydroxy-5beta-cholest-24-enoyl-CoA[x] + H2O[x] => (24R,25R)3alpha,7alpha,24-trihydroxy-5beta-cholestanoyl-CoA[x] |
| MAR08561 | 20-OH-LTB4[r] + O2[r] => 20-COOH-LTB4[r] + H+[r] + H2O[r] |
| MAR07701 | H+[c] + NADPH[c] + O2[c] => H2O2[c] + NADP+[c] |
| MAR01575 | 20-OH-LTB4[r] + NADPH[r] + 1.5 O2[r] => 20-COOH-LTB4[r] + 2 H2O[r] + NADP+[r] |
| MAR04423 | H2O[e] + O2[e] + putrescine[e] => 4-aminobutanal[e] + H+[e] + H2O2[e] + NH3[e] |
| MAR08606 | H2O[c] + O2[c] + putrescine[c] => 4-aminobutanal[c] + H+[c] + H2O2[c] + NH3[c] |
| MAR00059 | 3alpha,7alpha-dihydroxy-5beta-cholest-24-enoyl-CoA[x] + FADH2[x] + 0.5 O2[x] => (24R,25R)3alpha,7alpha,24-trihydroxy-5beta-cholestanoyl-CoA[x] + FAD[x] |
| MAR07703 | glycolate[c] + O2[c] => glyoxalate[c] + H2O2[c] |
| MAR07706 | glycolate[x] + O2[x] => glyoxalate[x] + H2O2[x] |
| MAR06539 | inositol[c] + O2[c] => glucuronate[c] + H+[c] + H2O[c] |
| MAR06606 | ascorbate[c] + urate radical[c] <=> monodehydroascorbate[c] + urate[c] |
| MAR06611 | H2O[c] + O2[c] + urate[c] <=> 5-hydroxyisourate[c] + H2O2[c] |
| MAR08017 | H2O[x] + lysine[x] + O2[x] => 6-amino-2-oxohexanoate[x] + H+[x] + H2O2[x] + NH3[x] |
| MAR08021 | L-pipecolate[x] + O2[x] => 1-piperideine-6-carboxylate[x] + H+[x] + H2O2[x] |
| MAR05390 | H2O[c] + methionine[c] + O2[c] => 4-methylthio-2-oxobutanoic acid[c] + H+[c] + H2O2[c] + NH3[c] |
| MAR06770 | H2O[c] + O2[c] + tyrosine[c] => 4-hydroxyphenylpyruvate[c] + H+[c] + H2O2[c] + NH3[c] |
| MAR07647 | D-alanine[x] + H2O[x] + O2[x] => H+[x] + H2O2[x] + NH3[x] + pyruvate[x] |

[*] This reaction is a variant of another reaction in the model, with the only difference that this reaction generates ROS. It was removed for convenience; it is not needed in the model since we don’t specifically model ROS.

### Supplementary notes

#### Supplementary note 1 – GECKO Light

GECKO Light is based on strategies previously developed and implemented in GECKO Toolbox[13,14], but with the aim to produce minimal models, similarly to what has been done in sMOMENT[15]. A metabolite "prot_pool" is added to the model, and each reaction associated with an enzyme cost consumes this metabolite. The stoichiometric coefficient *N* is calculated as

$$N = \frac{\sum_i M_{w,i}}{k_{cat}}$$

where $M_{w,i}$ is the molecular weight of part *i* in the complex, where non-complex enzymes are treated as a complex with just one part and $k_{cat}$ represents the catalytic efficiency of the enzyme. In cases where several isozymes convey the same reaction, the stoichiometric coefficient of the isozyme with the lowest protein cost is used.

Reactions with empty gene rules are assigned a standard protein cost (the median over all stoichiometric coefficients for "prot_pool" set in the model). Exchange reactions and transport reactions only involving a single metabolite are excluded, as well as reactions marked as spontaneous in Human1.

To hinder poorly matched outlier $k_{cat}$ values from dominating the simulation, GECKO Light fills in a standard protein cost for reactions with missing gene rules and corrects unrealistically low $k_{cat}$ values by not allowing values below a threshold (1 $s^{-1}$). This value is at the low end of the range of $k_{cat}$ values reported for most enzymes (1- 100 $s^{-1}$) (21506553) and is of the same order of magnitude as the $k_{cat}$ of the RuBisCO enzyme, known to be a particularly slow enzyme[16].

A constrained exchange reaction of the "prot_pool" metabolite (called "prot_pool_exchange") is also added to the model. The constraint on the total metabolic enzyme usage was fit to experimental values of 11 cell lines from NCI-60 for which the specific growth rate and metabolite uptake rates were deemed reliable[17–19], resulting in a value of 0.022 g/gDW.

### Supplementary note 2 – diffusion model

It is challenging to accurately model and quantify the maximum absolute fluxes of metabolites from the blood stream into the tumor. These fluxes depend on many parameters, such as capillary permeability and the geometry of the tumor and blood vessels. A more practical approach is to instead try to quantify the fluxes of metabolites relative to those of other metabolites, which can be useful for estimating which metabolites are potentially in short supply and thereby limiting for tumor growth. We assume a model where the influx of metabolites from the blood to the tumor is driven by diffusion.

The diffusion process is modeled axisymmetrically in two dimensions with radial flux out from the capillary wall, assuming that the capillary is long and that we therefore have no net flux along the capillary (Fig A). In steady state, we assume a fixed concentration of each metabolite just outside the capillary wall, which is proportional to its concentration in the blood ($c_b$). While it is hard to estimate the permeability of the blood vessel, such an assumption is reasonable under the assumptions that the blood vessel wall is thin, the concentration is constant in the blood vessel, and the metabolites are much smaller than the openings in the blood vessels. We then for simplicity assume an even distribution of cells in the tumor space, where the tumor space is divided into two regions for each metabolite. In tumor region A, closest to the capillary, there is an excess of the metabolite and cells only take up some of the available metabolite influx. In tumor region B, further away from the capillary, cells are starving and consume all the available metabolite influx. It is also assumed that the proportions of different cell types do not vary across the volume, and that cells operate according to the same metabolic regime regardless of distance to the capillary. The latter means that the uptake $u_i$ of metabolite $i$ at any distance from the capillary can be expressed as

$$u_i = g u_{r,i}$$

where g is a proportionality constant and $u_{r,i}$ is the uptake of metabolite $i$ at an arbitrarily chosen distance from the capillary that is the same for all metabolites. The border between region A and B is at different distances from the capillary for different metabolites. It is however important to realize that the metabolites of interest are the growth-limiting metabolites, since the estimate of the upper bounds of other metabolites have no effect on growth. In practice, all limiting metabolites are mainly used for ATP production. To simplify the model, we assume that the border between the regions A and B is at the same place for those metabolites, although this assumption may not hold for all metabolites. To simplify the calculations further, we use the approximation that this border is at the same distance from the capillary ($x_b$) for all metabolites, since the upper bounds of non-limiting metabolites has little effect on the modeling results.

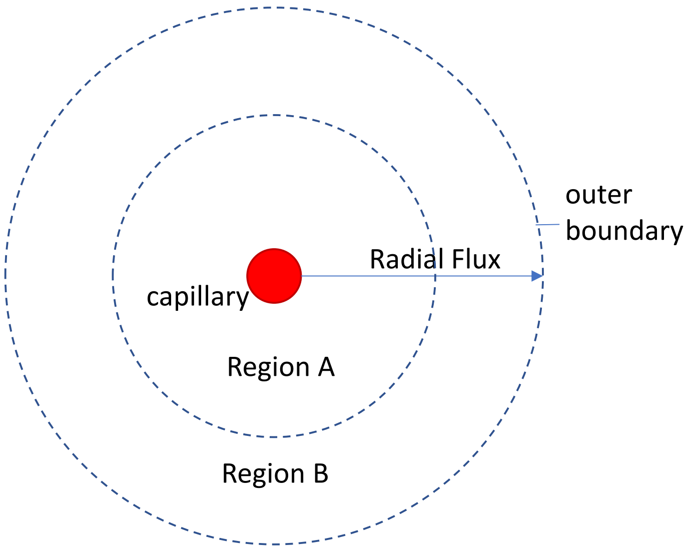


*Fig. A. The diffusion model.*

Diffusion is governed by Fick's first and second law, under the assumption that we view the tumor as an admixture in a medium, where the metabolite concentrations and gradients are "small" (which is the case here, the concentration is on the scale of around 5,000 µM at the most (for glucose) and the gradients are investigated at steady state conditions, giving no sharp concentration "edges"). The metabolite flux is determined by Fick's first law:

$$\boldsymbol{J}(\boldsymbol{x}) = -D\nabla \mathrm{c}(\boldsymbol{x}) \quad (1)$$

where **J** is the flux, *D* the diffusion coefficient, and $\nabla c(\mathbf{x})$ is the metabolite concentration gradient in three-dimensional space (where **x** normally is a three-dimensional vector, in our model a two-dimensional vector). We can determine $\nabla c(\mathbf{x})$ from boundary conditions and Fick's second law:

$$\frac{\partial c(x,t)}{\partial t} = \nabla \cdot (D\nabla c(\boldsymbol{x}, t)) \quad (2)$$

where *t* is time and $\nabla\cdot$ is the divergence of the vector field $D\nabla c(\mathbf{x}, t)$. For the problem at hand, we focus on steady state conditions, meaning that the left-hand side of Eq. 2 can be reduced to a sink which is constant over time (metabolite uptake) and that the concentration gradient is not dependent on time. We move the left-hand side to the right of Eq. 2. In Region A, the uptake rate is constant, and can simply be modeled as $-A$, where $A$ is a constant. In region B, we assume that cells take up a proportion of the metabolites that diffuse to the cell surface (with the same proportionality constant). This can be modeled as $-BDc(\mathbf{x})$, where $B$ is a constant. This assumes that for many substrates, such as glucose and amino acids, the most effective transporters have similar $K_m$ values. Since we assume that cells behave optimally, they will balance the expression of these transporters to obtain a similar transport capacity for all limiting metabolites, which will give a similar value for the constant $B$ for those metabolites.

If the cells in region A are assumed to behave the same way as cells on the border between region A and region B, we can assume that the metabolite uptake is the same for those cells. We can then replace the sink term $-A$ with $-BDc(\mathbf{x}_b)$, where $x_b$ represents the radial position of the border.

For simplicity, we assume that the diffusion constant *D* doesn't vary within the tumor, which means that D can be moved out of the divergence. We can then rewrite Eq. 2 as

$$0 = \nabla \cdot \left(D\nabla c(\boldsymbol{x})\right) - \frac{\partial c(x,t)}{\partial t} = D\nabla \cdot \left(\nabla c(\boldsymbol{x})\right) - \frac{\partial c(x,t)}{\partial t} = D\Delta c(\boldsymbol{x}) - \frac{\partial c(x,t)}{\partial t} \quad (3)$$

where $\Delta$ is the Laplace operator. We now insert the sink term for Region A:

$$0 = D\Delta c(\boldsymbol{x}) - A = D\Delta c(\boldsymbol{x}) - BDc(\boldsymbol{x_b}) \quad (4)$$

from which the diffusion constant can be removed:

$$0 = \Delta c(\boldsymbol{x}) - Bc(\boldsymbol{x_b}) \quad (5)$$

Similarly, we add the terms for Region B:

$0 = D\Delta c(\boldsymbol{x}) - BDc(\boldsymbol{x})$ (6), from which the diffusion constant can be removed:

$$0 = \Delta c(\boldsymbol{x}) - Bc(\boldsymbol{x}) \quad (7)$$

Given our assumptions, the concentration gradients for the limiting metabolites are not dependent on their diffusion coefficients in any of the regions.

To determine the concentration gradient from this equation at maximum metabolite uptake, we need boundary conditions. In our case, Dirichlet boundary conditions are suitable since they specify concentrations, not fluxes, and thereby do not involve the diffusion coefficient. We assume that in steady state there is a fixed concentration of the metabolite just outside the capillary wall, which is proportional to the concentration in the blood ($c_b$). It is hard to estimate the permeability of the blood vessel, but under the assumption that the blood vessel wall is thin, that the concentration is constant in the blood vessel, and that the metabolites are much smaller than the openings in the blood vessels, this is a reasonable assumption.

The second boundary condition used is that at infinite distance from the blood vessel, the concentration is zero. This is a commonly applied assumption that is required to be able to solve the concentration equation.

To solve the problem, we would need additional data, such as physical distances, etc., which is difficult to acquire. It is however not necessary to solve the problem to estimate the relative maximal fluxes of metabolites – we can directly draw some conclusions from the model given the assumptions: 1) The concentration gradient of a limiting metabolite is not dependent on the diffusion coefficient, since both Eq. 5 and 7 and the boundary conditions are independent of the diffusion coefficient. This means that the gradient $\nabla c(\boldsymbol{x})$ is approximately the same for all limiting metabolites, given the same concentration in the blood. 2) The concentration gradient $\nabla c(\boldsymbol{x})$ of a limiting metabolite is approximately proportional to the metabolite concentration in the blood. This follows from that at each boundary condition, the remaining concentration for the limiting metabolites is proportional to the concentration in blood (zero far out and proportional to the blood near the blood vessel), which intuitively means that the concentration gradient divided by the concentration in blood is approximately the same for all limiting metabolites.

Applying Fick's first law, we can thereby model the maximum influx of a limiting metabolite into any point of the tumor as

$$J(\boldsymbol{x}) = -D\nabla c(\boldsymbol{x}) = aDc_b \quad (8)$$

where the proportionality constant *a* is the same for all metabolites. For metabolic modeling purposes, the uptake flux of each metabolite can thus be constrained to this value. A low value of *a* represents the situation at a large distance from the capillary, while a high value gives conditions closer to the blood vessel. This equation can be generalized to

$$U_i = aD_i c_{b,i} \quad (9)$$

where $U_i$ is the estimated upper bound for the uptake flux of metabolite *i*, $D_i$ is the diffusion coefficient for metabolite *i*, and $c_{b,i}$ is the concentration of metabolite *i* in the blood.

There are many uncertainties that will introduce errors in the diffusion model. The model is based on four major assumptions: 1) The border between region A and region B is at the same place for all metabolites. 2) The cells have the same relative uptake rate of the growth-limiting metabolites regardless of the value of *a*, which is a simplification of the complex behavior of cells; 3) The cells are equally good at taking up different metabolites; and 4) We assume that the blood is constantly replenished and that the concentration of metabolites and oxygen in the blood are unaffected by cellular uptake.

Assumption 1 is likely violated for the limiting metabolites, which can be suspected from the FVA presented in Fig. S3. The violation is not expected to have a large impact on the modeling results but may introduce small errors in the estimated uptake bounds for some metabolites. The error manifests as a small under- or overestimation of the sink terms in Eq. 4 and Eq. 6 for some metabolites, but these errors are deemed small compared to the differences in metabolite concentrations in blood between metabolites.

Assumption 2 is likely violated to a certain extent since different absolute metabolite concentrations are expected to make the cells operate according to different metabolic regimes, which will introduce errors to the estimated uptake constraints. An example of this is the Warburg effect, where cells closer to the capillary are expected to take up a lower fraction of oxygen (compared to other metabolites) than cells further away from the blood vessel.

The third major assumption, where we assume that the cell is equally good at taking up different metabolites, is based on that most metabolites are imported to cells via effective transporters with low $K_m$ values. An example is the glucose transporter GLUT1, $K_m$ = 1-2 mM for glucose [20], and the glutamine transporter SNAT1, with reported $K_m$ = 0.23-0.30 mM [21]. It is assumed that differences in $K_m$ and $k_{cat}$ between the transporters can be compensated for by different abundancies of the transporters in the cell membrane.  Oxygen is an exception, which behaves very differently from the metabolites. Oxygen easily diffuses into cells through the membrane; the diffusion through the cell membrane is reported to be negligibly lower than in water [22]. Furthermore, the diffusion coefficient of oxygen is much higher than for the metabolites, making it more sensitive to errors in the diffusion model. Regardless of these differences, we still modeled oxygen uptake the same way as for the metabolites, although the relative constraint for oxygen uptake likely is less reliable.

Assumption 4, where the blood is expected to be constantly replenished with nutrients and thereby have constant metabolite concentrations throughout the tumor, may not hold. Depletion of oxygen in blood vessels is a well-known phenomenon referred to as acute hypoxia [23], and we can assume that the same effect to a certain extent is available for other metabolites. As long as the limiting metabolites are depleted in an equal way that is not a problem. However, oxygen behaves differently since the concentration is replenished from hemoglobin in the blood, and metabolites with higher diffusion coefficients may be more depleted in the blood. While the differences between most metabolites likely are small, oxygen may stand out together with lipids and albumin. To investigate the effect of violations of this assumption, we also modeled the metabolism using a blood flow model, where the metabolite uptake bounds are limited by blood flow and not diffusion (Note S3).

There are also uncertainties in the diffusion coefficients. For example, the diffusion constant of lipids and albumin are very uncertain, since they may be physically hindered to move in the TME due to their size. However, between metabolites such as glucose and amino acids, the difference in the diffusion coefficients is modest, and measurement errors there are not expected to have a large impact on the modeling results.

Taking all error sources into consideration, the uptake constraints are to be seen as an approximation of the true value. The uptake constraints for oxygen and lipids are expected to be more uncertain than the rest. Oxygen levels are however not that critical for the modeling results, we only need the oxygen levels to be low, which is a well-known fact in tumors. Results regarding lipids, however, are more uncertain and should be interpreted with care. In general, the diffusion model is also more reliable for higher values of a, since a smaller part of the regions where cells take up metabolites have been passed, reducing the errors introduced by some of the assumptions.

### Supplementary Note 3 – blood flow model

The diffusion model operates under the assumption that the metabolite availability for a certain cell is largely determined by its distance to the closest blood vessels, assuming that the blood supply in those blood vessels is sufficient. In such a case, the concentrations of metabolites available for diffusion into the tumor can be estimated to standard concentrations in blood. However, there is evidence that blood vessels in tumors are abnormal, leading to poor blood flow into the tumor [24]. There is therefore a risk of metabolite depletion in the blood and it could be that the blood flow rather than diffusion is the major limiting factor for metabolite supply to the cells. As an alternative to the diffusion model, we therefore also investigated the optimal metabolism using a blood flow model for limiting metabolite uptake rates.

The influx of metabolites into the tumor is assumed to be proportional to that of the standard blood concentrations. In the blood flow model, we assume that all cells get access to a proportion of the metabolites flowing into the tumor. That proportion varies across cells, where some regions have a higher blood flow than others. In the blood flow model, we thus limit the upper uptake bound $U_i$ of each metabolite *i* to

$$U_i = ac_{b,i} \quad (10)$$

where *a* is a proportionality constant (just as in the diffusion model) and $c_{b,i}$ is the standard concentration in blood of metabolite *i*. While it could be argued that the diffusion is still in play, the diffusion works in two ways. Given the same concentrations, metabolites with a higher diffusion coefficient can diffuse to the cells at a higher flux but will also be depleted faster in the blood, thereby lowering the concentration and reducing the effect. We have therefore chosen not to include the diffusion coefficient in the blood flow model.

A difference in the blood flow model compared to the diffusion model is the concentration of oxygen used. In the blood flow model, we assume that most of the oxygen will eventually be exported out of the blood vessel regardless of if it is bound to hemoglobin or not, and we therefore use the total oxygen consumption, estimated to 9,200 µM [4], which increases the oxygen availability substantially compared to the diffusion model case.

### Supplementary Note 4 – metabolite concentrations in blood

Since the diffusion model requires concentrations of metabolites in blood, we retrieved blood plasma measurements from several sources to form a collection of 69 metabolites with associated concentration values. The metabolites were filtered on the criteria that the metabolite must be represented in the model and have a blood plasma concentration above 1 µM. Specifically, we downloaded the average concentration of 94 polar metabolites in blood plasma from 8,413 healthy patients in a Japanese cohort [5] , of which 64 passed the filtering criteria, which also included removal of lipids. In addition to these metabolites, we also allowed for uptake of albumin, since cancer cells are known to take up albumin as a source of amino acids, at a plasma mass concentration of 40 g/l [25].

Due to the high number of lipid species we grouped the lipids into two categories: sterols and other lipids, where the latter included fatty acyls, glycerolipids, glycerophospholipids, and sphingolipids. Sterols were represented by cholesterol in the model, while the other lipids were assumed to eventually be converted to a composition of fatty acids defined by the metabolite "NEFA blood pool in", which is predefined in Human1 to match the composition in blood plasma. Lipid concentrations in blood plasma were downloaded from literature[26] and summed up to form mass concentrations in blood plasma for our two lipid groups. The average molecular weight of the "NEFA blood pool in" was calculated from its composition, and the mass concentrations were then converted to blood plasma concentrations for cholesterol and "NEFA blood pool in".

Estimation of the oxygen concentration is a special case since the concentration for the diffusion model should be that of free oxygen and not include oxygen bound to hemoglobin. The free oxygen has been estimated to only be around 1-2 % of the total oxygen in blood, at a concentration of around 100 µM [4]. In addition, any remaining metabolites that are essential for growth were added either unbounded (for metabolites that do not actively contribute to growth, such as water etc.) or with an upper bound of ten times the lowest upper bound present for the other metabolites, which is a level tested to be sufficient but not high enough to have a large impact on growth.

To test the validity of the metabolite concentration values, we downloaded and curated blood plasma concentration values from the human metabolome database (v. 4.0), HMDB[1,2], which contains concentration data from a large collection of studies. The metabolite concentrations in our metabolite collection were in good agreement with HMDB for the metabolites present in both sources (Fig. S1). In addition, we scanned HMDB for metabolites that were not present in our metabolite collection, yielding a collection of additional metabolite candidates (Table S1) and a list of metabolites that could not be mapped to the model (Table S2), both only containing metabolites with modest concentrations. To reduce the complexity of the model we decided not to include any of the new candidates since they were not deemed important and concluded that none of the unmapped metabolites would have a large effect on our modeling outcome. Our collection of metabolites with associated concentrations was hence deemed suitable for use with our model. The metabolite concentrations are available in Table S3.

#### Supplementary Note 5 – metabolite diffusion coefficients in tumors

The diffusion model requires the diffusion coefficients for the metabolites in the model. Reliable diffusion coefficients for the tumor microenvironment are scarce or unavailable, and it is difficult to know how diffusion coefficients differ between different fluids. Some values are available for blood plasma[27], although these were based on nuclear magnetic resonance (NMR) measured in two dimensions, which has been reported to be less reliable[28]. We settled for using diffusion coefficients measured in mouse seminal fluid for 16 of the polar metabolites in our metabolite collection[28]. We predicted the diffusion coefficients for the remaining small polar metabolites based on a linear model based on molecular mass (Fig. S2), which is reasonable as long as the masses are within a reasonably small range[29]. The diffusion coefficients for albumin[30] and oxygen[31] were collected separately from the literature. Lipids are not soluble in water and are therefore transported together with albumin or a lipoprotein, or potentially as droplets. These are large particles that do not diffuse well, and we modeled the diffusion of these particles by using the diffusion coefficient of albumin for all lipids. The metabolite concentrations and diffusion coefficients are available in Table S3.

### Supplementary Note 6 – Glutamine addiction and the NADH/$FADH_2$ ratio

Amino acid metabolism in the genome-scale model is tightly connected to the effects of NADH/$FADH_2$ production in the model. In normal conditions, these substrates can be oxidized by the electron transport chain to yield ATP. However, for fast growing cells with sufficient access to nutrients and oxygen, enzymatic capacity becomes limiting for growth and the model compensates by reducing its use of the ETC. The ETC complexes, of particular interest complex I, are large and slow and there are other strategies that can yield more ATP per enzyme usage. While aerobic glycolysis is the main alternative for producing ATP, the TCA cycle also remains an option, but can only be used if the NADH/$FADH_2$ generated in the cycle can be oxidized. Thus, if the cell can either generate less NADH/$FADH_2$ while running the TCA cycle or use alternative pathways to oxidize these substrates, the flux through the TCA cycle can be increased, contributing to the total ATP production. The NADH/$FADH_2$ ratio is also important, since NADH requires the use of complex I while $FADH_2$ does not, and the enzymatic cost per ATP produced is higher if complex I is used. However, the oxidation of NADH/$FADH_2$ by the ETC under hypoxic, nutrient-deprived conditions is limited by oxygen availability rather than enzyme usage, so it becomes important to get as much ATP as possible out of each oxygen molecule. Since the oxidation of $FADH_2$ is coupled to the reduction of ubiquinone to ubiquinol without generating any proton motive force, the ratio of NADH to $FADH_2$ processed by the ETC should be as high as possible. Depletion of NADH under such conditions may therefore have a negative effect on ATP production, and it may be beneficial to generate NADH by other means than running the TCA cycle to avoid generation of $FADH_2$.

While we do not predict such needs with the model, another important factor in some hypoxic conditions may be the ability of the TCA cycle to produce building blocks for the cell. For example, when cell lines are grown in severe hypoxic conditions with full access to a nutrient-rich medium, ATP could to a large extent be generated through glycolysis, and ATP generation may not be critical. The possibility to dispose of NADH to enable flux through the TCA cycle for the purpose of generating TCA cycle intermediates such as α-ketoglutarate as building blocks for the cell may in such cases be an important function in the cell[32].

As mentioned in the main text, glutamine can be used as substrate to the TCA cycle instead of pyruvate to increase the flux through the TCA cycle. This leads to less NADH production per round in the TCA cycle but requires disposal of aspartate. Aspartate can be exported directly but can also be converted to lactate (via fumarate, malate, and pyruvate) without requiring ATP or altering the redox balance, which may explain why aspartate export is not observed. This process couples the oxidation of NADH to $NAD^+$ with the reduction of $NADP^+$ to NADPH, which may impact other processes. In addition, export of dihydroorotate has also been reported in hypoxic conditions, posing as an alternative pathway for disposing of aspartate[33]. Interestingly, it has been reported that different cell lines have different strategies for aspartate supply; some do not express aspartate transporters and rely on ETC activity for aspartate production, which is consistent with our modeling results, while others rely on import[34]. An additional benefit of using glutamine as substrate for the TCA cycle instead of pyruvate is that less ROS is produced, since complex I is run less and complex I produces substantially more ROS than complex II[35]. Another possibility is that glutamine can be converted to α-ketoglutarate and further via the TCA cycle in reverse to acetyl-CoA to support lipogenesis, which has been reported to be an important pathway in cell lines cultured under hypoxia[36]. While this is not predicted by the model since many lipids are directly supplied by the diffusion model, such a behavior presents another possibility to dispose of NADH in hypoxia using glutamine.
